# A ripening-associated ethylene response factor ERF.D7 activates ARF2 orthologs to regulate tomato fruit ripening

**DOI:** 10.1101/2022.07.11.499599

**Authors:** Priya Gambhir, Adwaita Prasad Parida, Vijendra Singh, Utkarsh Raghuvanshi, Rahul Kumar, Arun Kumar Sharma

**Affiliations:** Department of Plant Molecular Biology, University of Delhi South Campus, New Delhi 110021, India; Department of Plant Sciences, School of Life Sciences, University of Hyderabad, Hyderabad 500046, India

**Keywords:** ARF, auxin-ethylene crosstalk, carotenoid accumulation, ERF, fruit ripening, fruit quality, hormone signalling, overexpression, RNA interference, tomato

## Abstract

Despite the obligatory nature of ethylene to climacteric fruit ripening and the identification of 77 Ethylene Response Factors (ERFs) in the tomato genome, the role of only limited ERFs has been validated in the ripening process. Here, using a comprehensive morpho-physiological, molecular and biochemical approach, we demonstrate the regulatory role of SlERF.D7 in tomato fruit ripening. The expression of *SlERF.D7* positively responds to exogenous ethylene and auxin treatments, most likely in a RIN-independent manner. Overexpression of *SlERF.*D7 promotes ripening and its silencing has the opposite effects. Alterations in its expression modulated ethylene production, pigment accumulation, and fruit firmness. Consistently, genes involved in ethylene biosynthesis and signalling, lycopene biosynthesis, and cell wall loosening were upregulated in the overexpression lines and downregulated in RNAi lines. These transgenic lines also accumulated altered levels of IAA at late-breaker stages. A positive correlation between *ARF2* orthologs transcripts and *SlERF.D7* mRNA levels and the fact that *ARF2A* and *ARF2B* are direct targets of SlERF.D7 underpinned the perturbed auxin-ethylene crosstalk for the altered ripening program observed in the transgenic fruits. Overall, this study uncovers that SlERF.D7 positively regulates SlARF2A/B abundance to amalgamate auxin and ethylene signalling pathways for controlling tomato fruit ripening.

**Highlight:** The present study identifies a new ripening regulator, SlERF.D7, that controls tomato fruit ripening by activating other known-ripening regulators, *SlARF2A* and *SlARF2B*.

## Introduction

Owing to the agronomical position tomato upholds, a comprehensive dissection of its intrinsic ripening program is imperative to unravel the molecular mechanisms governing vital qualitative and quantitative fruit-related traits. The original concept of the fruit maturation process in climacteric fruits is based on a classic linear prototype guided by the precise spatio-temporal expression of the genes of ethylene biosynthesis and signalling (Alba *et al*., 2005; Kumar *et al*., 2016; Sravankumar *et al*., 2018). However, zooming in on the transcription networks activated during ripening in fleshy fruits has revealed a well-defined information system in which a multitude of hidden layers tightly control the ethylene biosynthesis and signalling to produce ripened fruits as an output. One such regulatory circuit involves crosstalk between ethylene and auxin, in which the latter antagonistically targets major facets of ethylene-regulated fruit ripening (Böttcher *et al*., 2010; Schaffer *et al*., 2013; Sravankumar *et al*., 2018). Auxin and ethylene are two cornerstones of overall fruit development and are assumed to be involved in inevitable trade-offs. i.e., the ability of ethylene to trigger fruit ripening occurs at the expense of disruption of auxin biosynthetic and signalling machinery (Given *et al*., 1988; Zaharah *et al*., 2012; Kumar *et al*., 2014; Sravankumar *et al*., 2018). Experimental validation of this trade-off was reported when the exogeneous application of auxin inhibitor, p-chlorophenoxyisobutyric acid, to tomato fruits mimicked ACC treatment and displayed early signs of fruit ripening (Su *et al*., 2015).

Deepening insights into the connection between signal transduction components of auxin and ethylene roots back to the era of identification of ethylene insensitive mutants with defects in auxin transporters, *aux1* and *ethylene insensitive root1/ pinformed 2* (*eir1/pin2*) (Pickett *et al*., 1990; Luschnig *et al*., 1998). These mutants exhibited root growth inhibition synergistic with the effect of auxin on this process (Rahman *et al*., 2001; Swarup *et al*., 2002). Apart from being collaborators, ethylene and auxin have exemplified competitiveness in lateral root initiation in *Arabidopsis* whereby auxin have been shown to promote lateral root formation, and elongation with ethylene negatively regulated the primary as well as lateral root elongation (Negi *et al*., 2008, 2010; Ivanchenko *et al*., 2008; Muday *et al*., 2012). The centrality of auxins in defining regions of meristem growth in roots and shoots has long been recognized (Benková *et al*., 2003; Blilou *et al*., 2005). What is fascinating is that the pattern of auxin transport in these regions is strongly influenced by ethylene (Růžička *et al*., 2007). Once accumulated, auxin then initiates the repression of ethylene mediated root growth phenotype (Lewis *et al*., 2011). Additionally, the points of convergence between auxin and ethylene at the transcriptional level have been extensively studied due to documentation of their receptor and signalling mutants in the public domain. Ethylene-insensitive root growth phenotypes were observed in auxin receptor mutant (*TIR1*), auxin transport mutants (*AUX1* and *EIR/AGR/PIN2*) as well as in mutants deficient in auxin response (*AXR2/IAA7* and *AXR3/IAA17*) (Pickett *et al*., 1990*a*; Luschnig *et al*., 1998; Stepanova *et al*., 2005; Muday *et al*., 2012). In parallel, ethylene receptors are up-regulated in fruits by auxins (Gillaspy *et al*., 1993; Jones *et al*., 2002; Trainotti *et al*., 2007). These points of intersection serve two separate and important roles: regulating global plant architecture and conferring local robustness on development.

In climacteric fruits, an absolute requirement of increased ethylene production via the upregulation of 1-aminocyclopropane-1-carboxylate synthase 2 (*ACS2)* and 1-aminocyclopropane-1-carboxylate oxidase 1 (*ACO1)* transcripts, coordinated with low auxin levels sets the stage for ripening (Alba *et al*., 2005; SravanKumar *et al*., 2018). Interestingly, mRNA levels of *ACS2*, *ACS4,* and *ACO1* are upregulated by auxin in tomato and peach (Jones *et al*., 2002; Trainotti *et al*., 2007). The physiological effects of both the hormones, at the genetic level, are brought about by their main transcriptional regulators, such as EIN3-like proteins (EILs) and ethylene response factors (ERFs) in the case of ethylene and auxin response factors (ARFs), AUX/IAA and Topless proteins in case of auxin. Genome-wide identification studies have revealed that the tomato genome harbors 77 ERF (Sharma *et al*., 2010; Pireello *et al*. 2012; Srivastava *et al*., 2019) and 22 ARF members. Accumulated evidence shows that ethylene controls the accumulation of some ARF transcripts during tomato fruit development signifying the possibility of involvement of auxin in climacteric fruit ripening (Jones *et al*., 2002). Likewise, several ERF genes are regulated by auxin (Trainotti *et al*., 2007; Pirrello *et al*., 2012). This two-way communication channel is feasible due to the multi-member gene families of ERFs and ARFs and the presence of both auxin and ethylene *cis*-regulatory elements in the promoter regions of these two sets of transcription factors (Muday *et al*., 2012; Zouine *et al*., 2014). Consistent with antagonism to ethylene, tomato fruit firmness and sugar metabolism have been reported to be partly regulated by *SlARF4* (Jones *et al*., 2002; Sagar *et al*., 2013). More recently, another auxin response factor, *SlARF2* has been proven to be a quintessential component of the regulatory network controlling fruit ripening. Silencing of *SlARF2* brought about dramatic ripening defects with a concomitant reduction in ethylene production and down-regulation of key ripening regulators *RIN*, *NOR*, and *CNR* in tomato (Hao *et al*., 2015; Breitel *et al*., 2016*a*).

Several ripening-induced ERF genes have been implicated in the regulation of fruit ripening in tomato (Sharma *et al*., 2010; Liu *et al*., 2014, 2016). Overexpression of one such gene, *LeERF1,* resulted in constitutive ethylene response with accelerated fruit ripening and enhanced fruit softening (Li *et al*., 2007; Liu *et al*., 2015). Another key piece of evidence further substantiating the importance of ERFs in ripening has come from *SlERF6,* which integrates ethylene and carotenoid pathways in tomato (Lee *et al*., 2012; Liu *et al*., 2016). Due to the issue of proposed overlapping functions among ERF members, evidence for the involvement of another ERF, *SlERF.B3,* in controlling carotenoid accumulation and ethylene production was demonstrated using a dominant repressor strategy (Liu *et al*., 2014). Although the contribution of a few of the ripening-induced ERFs in tomato fruit ripening has been elucidated, the function of the majority of these genes largely remains undocumented. Moreover, due to the constitutive overexpression or silencing of these genes in earlier studies, pleiotropic phenotypes unrelated to fruit ripening have also been reported (Li *et al*., 2007; Liu *et al*., 2018). Previously, we have identified and reported several ripening-induced ERFs (Sharma *et al*., 2010; Kumar *et al*., 2012; Srivastava *et al*., 2019). We have also characterized a fruit ripening-specific *RIP1* promoter (Agarwal *et al*., 2017). In the present study, we report the functional validation of a yet to be described ripening-induced ERF gene, *SlERF.D7*, for its roles in the regulation of fruit ripening traits in tomato by silencing it under *RIP1* promoter. First, we report that *SlERF.D7* transcripts are inhibited during fruit ripening in *rin* and *nor* mutants and *in-house* RIN-silenced lines (Kumar *et al*., 2012). Transcript profiling using quantitative real-time PCR (qRT-PCR) revealed strong induction of *SlERF.D7* mRNA levels upon exogenous ethylene and auxin applications. Using a reverse genetic approach, we demonstrate that *SlERF.D7* plays a pivotal role in fruit ripening via directly modulating the expression of *SlARF2,* thereby serving as a critical point of intersection between ethylene and auxin signalling pathways. Fruit-specific silencing of *SlERF.D7* under *RIP1* promoter results in a severe reduction in ethylene production, fruit firmness, and pigment accumulation of the transgenic fruit, a phenotype that resembles the *SlARF2* down-regulation line fruits (Hao *et al*., 2015; Breitel *et al*., 2016*a*). In contrast, *RIP1* driven ripening-specific overexpression of *SlERF.D7* fastened the ripening progression and enhanced fruit lycopene levels. Thus, using a suite of morpho-physiological, biochemical, pharmacological, and molecular tools, we identify a new regulator of tomato fruit ripening and provide critical insight into the auxin-ethylene controlled molecular circuitry that controls the ripening traits in tomato.

## Materials and Methods

### Plant materials and growth conditions

Tomato (*Solanum lycopersicum* L.cv Pusa Ruby) wild-type, *35S:RIN-RNAi* transgenic lines in Pusa Ruby background generated in our lab (RIAI05 and RIAI06) (http://hdl.handle.net/10603/389794), Ailsa Crag wild-type and *rin* (accession no. LA1795, in an unknown background) and *nor* (Ailsa Craig background) fruit ripening defective mutants were grown under standard greenhouse conditions: 14-h-day/10-h-night cycle, 25°C/20°C day/night temperature, 60% relative air humidity, and 250 µmol m^−2^ s^−1^ intense luminosity. Fruit pericarp tissue samples were collected from different fruit development and ripening stages, as described earlier (Kumar *et al*., 2012). For the measurement of time to ripen, flowers were tagged at anthesis. The number of days counted from anthesis to the appearance of the first symptoms of ripening was designated as the number of days taken to reach the breaker (Br) stage. The mature green (MG) stage was fixed one day before the Br stage. All fruit stages were harvested in three biological replicates (each replicate was represented by a pool of several fruits from different plants for each genotype used in the present study). Upon harvest, fruits were snap-frozen in liquid nitrogen to avoid injury.

### Plasmid construction and VIGS assay

Tobacco rattle virus (TRV)- based *pTRV1* and *pTRV2* vectors were used for VIGS experiments (Liu *et al*., 2002). A 300-bp fragment of the coding region corresponding to *SlARF2A* and *SlARF2B* was retrieved by us using the VIGS tool of the SOL Genomics Network Database (https://vigs.solgenomics.net/). Each 300-bp fragment was then PCR amplified from *Solanum lycopersicum* cv Pusa Ruby cDNA and inserted in the *pTRV2* vector. For double knock-down of *SlARF2A* and *SlARF2B*, a chimeric construct with fragments corresponding to both the genes was cloned and ligated into the same *pTRV2* vector. The primers used for VIGS assay cloning are listed in the Supporting table: Table S1. The empty *pTRV* vector was used as a control in the VIGS assays. The *pTRV::ARF2A*, *pTRV::ARF2B,* and *pTRV::ARF2AB* plasmids were verified by sequencing and mobilized into *Agrobacterium tumefaciens* strain GV3101. As described previously, VIGS was carried out on mature green fruits (35DPA) (Fu *et al*., 2005).

### Subcellular localization of SlERF.D7

The coding sequence (CDS) of *SlERF.D7* was cloned in frame with GFP reported gene into *pSITE-2CA* vector (primers are listed in Supporting table: Table S1). The empty *pSITE-2CA* vector was used as a control. *SlERF.D7-pSITE-2CA* and the control vectors were transferred to *A*. *tumefaciens* strain GV3101 and injected into 4-week-old tobacco leaves as described previously (Martin *et al*., 2009). GFP fluorescence was observed and captured by a laser confocal microscope (Leica TCS SP8, Germany) after 48 h of infiltration, as described previously by us (Kumar *et al*., 2015).

### *In silico* analysis of SlERF.D7, SlARF2A, and SlARF2B promoters

To identify putative *cis*-acting elements in the promoter sequences of *SlERF.D7, SlARF2A,* and *SlARF2B*, 2.5-kb upstream regions from the transcription start site of each gene were retrieved from the SOL Genomics Network database. After that, the sequences were scanned using the PLACE/signal search tool (http://www.dna.affrc.go.jp/PLACE/signalscan.html) to identify ethylene/auxin/fruit-specific/ripening-related *cis*-acting regulatory elements.

### Phytohormones treatment

For ethylene treatment to fruits, tomato fruits were harvested at the mature green stage of development and injected with a buffer solution containing 10 mM MES, pH 5.6, sorbitol (3% w/v), and 100 μM of Ethrel (2-Chloroethylphosphonic Acid, 40% Solution, Sisco Research Laboratories Pvt. Ltd. India). Similarly, auxin treatment to the mature green fruits were given by injecting a buffer solution 10 mM MES, pH 5.6, sorbitol (3% w/v) having 100 µM IAA. For inhibitors treatment, tomato fruits harvested at breaker fruits were infiltrated with the abovementioned buffer containing 100 µM 1-MCP and 100 µM PCIB, respectively. Briefly, similar-sized tomato fruits were injected using a 1 ml syringe with a 0.5 mm needle and inserted 3 to 4 mm into the fruit tissue through the stylar apex. The infiltration solution was gently injected into the fruit until the solution ran off the stylar apex and the hydathodes at the tip of the sepals. Only completely infiltrated fruits were used in the experiments. Control fruits were treated with the corresponding buffers only. After the treatment, fruits were incubated in a culture room at 26°C, under a 16 h light/8 h dark cycle with a light intensity of 100 μmol m^-2^s^-1^. After 24 h, the fruit pericarp was collected and frozen at -80°C until further use.

### RNA isolation and quantitative real-time polymerase chain reaction

Total RNA from the different tissues/organs/stages (cotyledon, root, leaf, stem, flower, and pericarp of fruits at IMG, MG, Br, Br+3, Br+5, and Br+10) of tomato plants was isolated using RNeasy Plant Mini Kit (Qiagen, Germany) by following manufacturer’s instructions. A provision for on-column DNase-treatment was provided in the kit. After the RNA isolation, 1-µg of total RNA for each sample was subjected to cDNA synthesis with high-capacity cDNA reverse transcription kit (Applied Biosystems, USA), according to the instructions provided in the manual. Gene-specific primers for qPCR were designed using PRIMER EXPRESS version 2.0 (PE Applied Biosystems, USA) with default parameters. Further, 2X Brilliant III SYBR® Green QPCR master mix (Agilent Technologies, USA) was used for a qRT-PCR reaction carried out in a Stratagene Mx3005P qPCR machine (Agilent Technologies, USA). Three independent RNA isolations along with three technical replicates were used for mRNA quantification. The expression values of genes were normalized using *GAPDH* and *ACTIN* gene expression values. Relative expression values were calculated by employing the 2^-ΔΔCT^ method (Livak and Schmittgen, 2001).

### Construction of *SlERF.D7* overexpression and silencing vectors and tomato transformation

To generate the *RIP1::SlERF.D7* OE transgenic plants, we amplified the full-length coding sequence and first cloned in pGEM-T Easy vector (Promega, USA) and finally mobilized into a binary vector, *pCAMBIA2300-RIP1-NOSt*, where we have already cloned the *RIP1* promoter. Similarly, to generate the *RIP1::SlERF.D7* RNAi transgenic plants, a 380 bp unique coding sequence of *SlERF.D7* was PCR amplified and cloned first in pUC19. The same fragment in sense and antisense orientation was cloned in juxtaposition, where an intronic sequence separated the two fragments. The assembled construct was finally mobilized into another binary vector *pBI121-RIP1-NOSt*. The cloning was confirmed by PCR, restriction digestion, and Sanger’s sequencing. The sequence-confirmed overexpression (OE) and silencing (RNAi) plasmids were mobilized from *E. coli* into *Agrobacterium* tumefaciens strain AGL1. After confirmation of this mobilization, the transformed *Agrobacterium* cultures were used to generate stable transgenic *RIP1::SlERF.D7* OE and RNAi lines in Pusa Ruby, as described previously (Maligeppagol *et al*., 2011). Primers used in constructing *RIP1::SlERF.D7* OE and RNAi constructs are listed in Supporting Table S1. The OE and RNAi lines were grown and analyzed for their morphological, biochemical, molecular, and physiological characterization in the T_2_ generation. The segregation analysis of kanamycin resistance in T_2_ progeny of *SlERF.D7 OE* and *SlERF.D7* RNAi transgenic tomato plants was done to obtain homozygous lines. Final experiments were carried out using homozygous lines from T_2_ or later generations.

### Fruit firmness measurement

The assessment of fruit firmness was carried out using TA.XT Plus Texture Analyser (Stable Micro Systems, England). Fruits that had undergone VIGS at the red ripe (Br+7) stage were subjected to a puncture test with a 2 mm needle probe, and the force and distance measurements were recorded. Fifteen fruits from each transgenic and tissue culture generated wild-type genotype were used in the stufy. Fruit firmness calculated is equivalent to the amount of force applied to penetrate the surface of the fruit. For fruit shelf-life measurement, ten fruits from the wild-type and transgenic lines were harvested at the MG stage and kept in the dark at room temperature, and photographed at regular intervals during the experiment.

### Ethylene measurement

Fruits at each developmental stage were harvested and placed in open 250-ml jars for 2 h to minimize the effect of wound-induced ethylene caused by the harvesting of the fruit. Jars were then sealed and incubated at room temperature for 4 h, and 1 ml of headspace gas was injected into Shimadzu QP-2010 Plus with Thermal Desorption System TD 20 (Shimadzu, Japan). Samples were compared with reagent-grade ethylene standards of known concentration and normalized for fruit weight. Ethylene in the headspace gas was measured thrice for each sample, with at least three biological samples for each ripening stage.

### Estimation of fruit pigments

Lycopene and β-carotenoid extractions for HPLC experiments were performed as described previously (Fantini *et al*., 2013). Briefly, 150 mg of ground lyophilized tomato fruit powder were extracted with chloroform and methanol (2:1 v/v); subsequently, 1 volume of 50 mM Tris buffer (pH 7.5, containing 1M NaCl) was added, and followed by the incubation of samples on ice for 20 min. After centrifugation (15,000 *g* for 10 min at 4°C), the organic phase was collected, and the aqueous phase was re-extracted with the same amount of chloroform. The combined organic phases were then dried by centrifugal evaporation and resuspended in 100 µl of ethyl acetate. A final volume of 20 µl was injected into a C-18 column in HPLC (Shimadzu, Japan) analysis. For each genotype, at least five independent extractions were performed.

### Estimation of titrable Acids

The amount of titrable acids (TA) was determined by titrating the fruit homogenates against 0.1 N NaOH solution using phenolphthalein as an indicator to the endpoint at pH 8.1 (Singh and Pal., 2008). The titrable acids were measured by using the following equation:

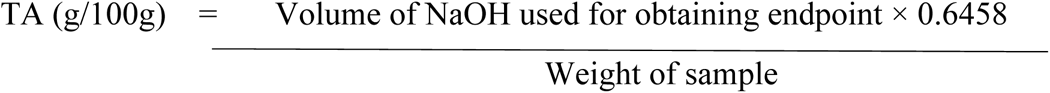

### Determination of total soluble solids

The quantification of total soluble acids (TSS) was done by measuring the refractive index of tomato juice through a portable refractometer (Erma Handheld Refractometer, India). The refractive index of water was initially calculated with the device to serve as zero error. A drop of the sample (tomato juice) of all the test samples was placed individually on the measuring surface beneath the Viewpoint Illuminator. Through the eyepiece, the readings were recorded at the point where the contrast line crossed the scale. The results have been expressed in degrees Brix (percentage of TSS in solution at 20°C).

The ripening Index for all the samples was calculated using the following formula:

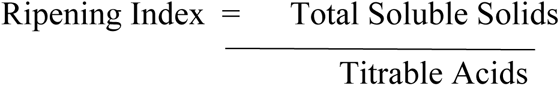

### Measurement of IAA in fruit pericarp tissue

Extraction and purification of IAA were carried out as previously reported with slight modifications (Edlund *et al*., 1995). Frozen samples were ground in liquid nitrogen, 50 mg fresh weight of the powdered sample was mixed with 1ml of 80% methanol containing 1% acetic acid (v/v), and 2 ng of [^13^C_6_]-IAA was added as an internal standard (Cambridge Isotope Laboratories, Andover, MA, USA). The sample was extracted for 2h at 4°C under continuous shaking. After centrifugation (10,000 *g*, 5 min), the supernatant was collected and concentrated in a vacuum. The sample was resuspended in 1 ml 0.01M HCl slurried for 10 min at 4 °C under continuous shaking with 15 mg Amberlite XAD-7HP (Organo, Tokyo, Japan). After removing the supernatant, the XAD-7HP was washed twice with 1% acetic acid. Samples were then extracted with CH_2_Cl_2_ (once with 400 µl and twice with 200 µl), and the combined CH_2_Cl_2_ fraction was passed through a 0.2 µm filter. After concentration in a vacuum, the sample was analyzed by GC-MS. Five independent samples were extracted and analyzed.

### Yeast-one-hybrid assay

For yeast-one-hybrid (Y1H) experiments, 1.5 kb long promoter sequences of *SlARF2A* and *SlARF2B* were amplified using tomato (*Solanum lycopersicum* L.cv Pusa Ruby) genomic DNA. The amplified products were cloned into the yeast expression vector *pHIS2*.1 and co-transformed with the *pGAD-T7-SlERF.D7* into yeast strain *Y187*. The binding of RIN to the promoter of *LeACS2* was taken as a positive control for this experiment. For negative control, the interaction of *SlARF3* promoter with *SlERF.D7* was used. The DNA-protein interaction was validated by transformant growth assays on SD/-Leu/-Trp/-His plates supplemented with 50 mM of 3-AT. Primers used in this section are listed in the Supporting table: Table S1.

### Transactivation of *SlARF2A* and *SlARF2B* promoters in *N. benthamiana*

To verify the DNA-protein interaction in planta, 1.5 kb upstream regions of *SlARF2A* and *SlARF2B* were used to drive the expression of GUS and GFP, and were designated as reporter vectors. The effector vector was constructed by cloning the full ORF of *SlERF.D7* in the binary vector *pBI121* driven by *CaMV35S* promoter. As previously described, both reporter and effector constructs were co-transformed into 4-week-old tobacco leaves GUS histochemical staining assay and GFP fluorescence measurement assay were done after 48 h of infiltrations. The NightSHADE LB 985 (Berthold Technologies USA) in vivo plant imaging system was employed to detect fluorescent signals with 5s exposure time. Data were analyzed by the IndiGO™ software. The average fluorescence signal for each sample was collected (cps, count per second). It was normalized using the tobacco leaves transformed with a reporter vector combined with the vector used as effector but lacking the ERF or ARF coding sequences. Three independent replicates were used for each analysis. Positive and negative controls were the same as reported earlier for the Y1H assay. Primers used in this experiment are listed in the Supporting table: Table S1.

### Electrophoretic mobility shift assay

The full-length *SlERF.D7* coding sequence was cloned in frame into pET-28a (to fuse in frame with 6X Histidne tag) for heterologous protein epression. The fusion protein construct was expressed in BL21 strain of *E.coli*. Recombinant SlERF.D7-6XHIS protein was purified by Ni^2+^gravity flow chromatography according to the manufactureer’s protocol (Ni-NTA Agarose, Qiagen) as described. For EMSA assay, the 50-bp probe covering the ERE element (AGCCGCC) derived from *SlARF2A* and *SlARF2B* promoter was labeled with digoxigenin as per the manufacturer’s protocol using DIG Oligonucleotide 3′-End Labeling Kit (Roche Diagnostics). The same unlabeled DNA fragment was used as a competitor. The binding reactions were performed at room temperature in binding buffer (10 mM Tris (pH7.5), 50 mM KCl, 1 mM DTT, 2.5% glycerol, 5 mM MgCl_2_, 0.5 mM EDTA, 50 ng /ml poly (dI-dC)) containing 1µg purified SlERF.D7-6XHIS fusion protein and 5ng probes. The reaction products were analyzed on 6% (w/v) native polyacrylamide gel electrophoresis. The products then transferred from the gel to Hybond N+ Nylon membrane (Amersham Biosciences) and detected using DIG Nucleic Acid Detection Kit (Roche Diagnostics) according to manufacturer’s protocol.

### GUS histochemical and activity assays

Histochemical staining of GUS in tobacco leaves was performed as described previously. Briefly, *N. benthamiana* infiltrated leaves were harvested and incubated in a substrate solution [50 mM sodium phosphate, pH 7.5, 0.5 mM K_3_Fe(CN)_6_, 0.5 mMK_4_Fe(CN)_6_, 1 mM X-gluc (5-bromo-4-chloro-3-indolyl-b-D-glucuronide)] at 37°C overnight. Tissues were then cleared in 70% ethanol overnight. For the fluorometric GUS assay, the infiltrated tobacco leaf discs were homogenized in 1 ml of extraction buffer (50 mM NaH_2_PO_4_, pH 7.0, 10 mM EDTA, 0.1% Triton X-100, 0.1% (w/v) sarcosyl (w/v), and 10 mM β-mercaptoethanol). The homogenate was centrifuged for 10 min at 12,000 *g* at 4°C, and 100 µl of the supernatant was mixed with 900 µl of assay buffer (1mM 4-methylumbelliferyl-D-glucuronide in the extraction buffer). The reaction mixture was incubated at 37°C for 1h and eventually stopped by adding a stop buffer (0.2 M sodium acetate). GUS activity was normalized to protein concentration in each crude extract and was calculated as pmole or nmole of 4-methylumbelliferone (4-MU) min^-1^mg^-1^protein. The Bradford method assessed the protein content using Bovine serum albumin (BSA) as a standard.

## Results

### Transcript profiling of *SlERF.D7* reveals a potential role in fruit ripening

First, to predict the function of SlERF.D7, its transcript abundance was assessed by quantitative qRT-PCR in different plant tissues/organs/stages, including cotyledons, leaf, stem, root, flower, and flower, different stages of tomato fruit development and ripening. The transcript levels of *SlERF.D7* were found to be low in vegetative organs. In contrast, it displayed a drastic up-regulation at the 5-days post breaker stage and reached its maximum levels at 10-days post breaker, indicating its prospective requirement in the ripening process (Fig. 1A). Next, we evaluated its mRNA abundance at different ripening stages, namely mature green (MG), breaker (Br), 3-days post breaker (Br+3), 5-days post breaker (Br+5), and 10-days post breaker (Br+10) in wild-type, RIAI05 and RIAI06 transgenic lines and same-age fruits of two ripening-mutants, *ripening-inhibitor* (*rin*) and *non-ripening* (*nor*) (Fig. 1B,C). Contrary to the wild-type fruits, the qRT-PCR analysis revealed no ripening-associated induction of *SlERF.D7* transcripts at B+5 and B+10 stages in *rin* mutant fruits (Fig. 1B). However, fruits from RIN suppressed transgenic lines (RIA105 and RIAI06), and *nor* mutant fruits exhibited a slight increase in the expression level of *SlERF.D7* at late-breaker stages, but the enhancement was significantly lower than the wild-type fruits. These results indicated the functional significance of the *SlERF.D7* gene during tomato fruit ripening.

**Figure 1.**
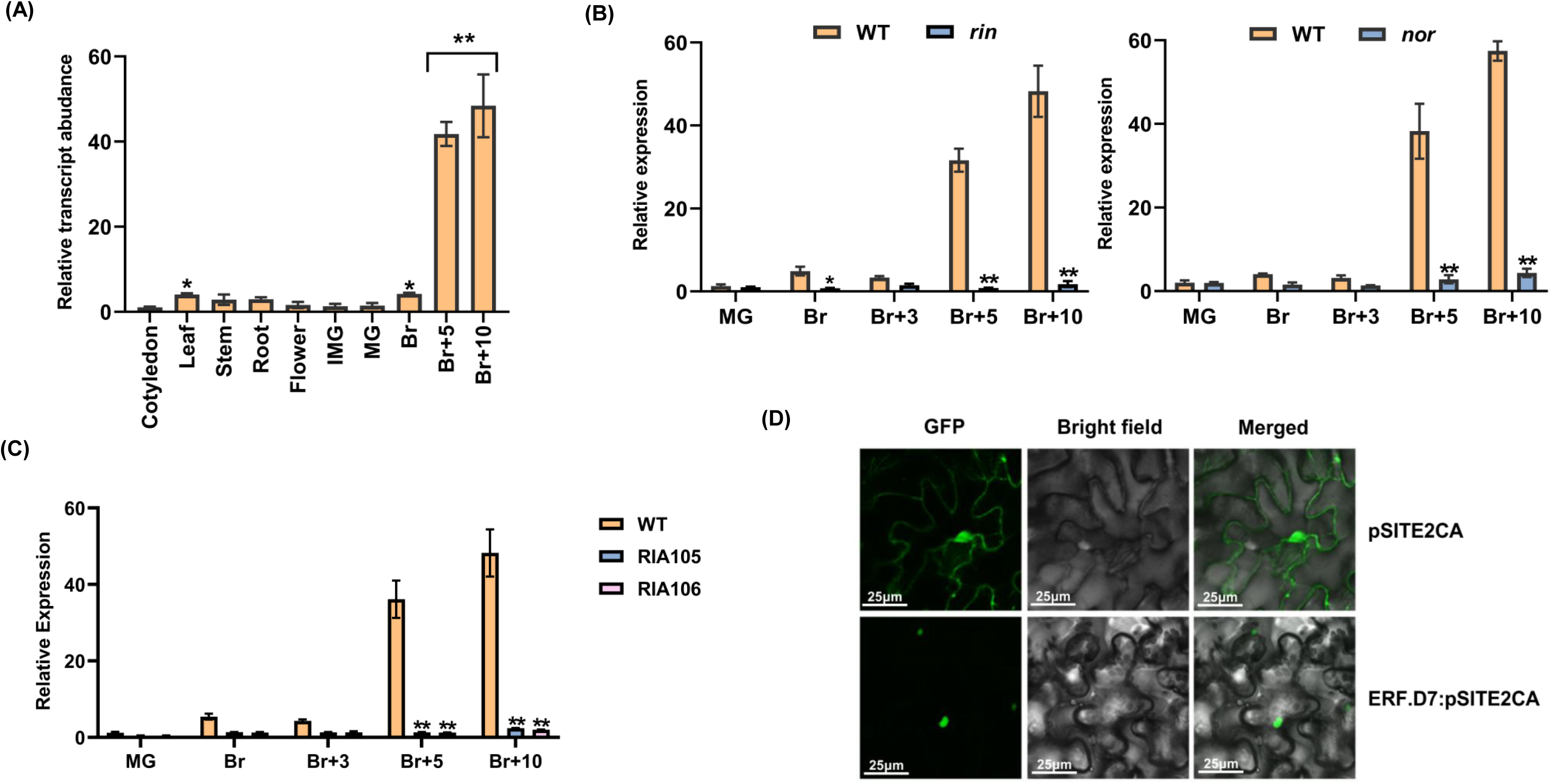
Transcript profiling of *SlERF.D7* (Solyc03g118190) and its subcellular localization. **(A)** Expression of *SlERF.D7* in various tissues, including cotyledon, roots, stems, leaves, flowers, and fruit at different developmental stages: immature green (IMG), mature green (MG), breaker (Br), 5-day after Br (Br+5) and 10-day after Br (Br+10) in wild-type (Pusa Ruby) . Values are means ± SD of three independent replicates. Asterisks indicate the statistical significance using ANOVA: *, 0.01 < P-value < 0.05; **, 0.001 < P-value < 0.01. **(B, C)** Relative expression levels (in fold change) of *SlERF.D7* in wild-type (Ailsa Craig), *ripening-inhibitor* (*rin*; accession no. LA1795, in the unknown background) mutant and *nor* mutant (in Ailsa Craig background) and 3S:RIN knockdown lines (RIA105 and RIA106; in Pusa Ruby background) at various stages of fruit ripening. Expression profiles were studied at different stages of fruit ripening by employing the qRT-PCR technique. The mRNA levels of *SlERF.D7* at the MG stage in WT were used as the reference for all stages. Values are means ± SD of three independent replicates. Asterisks indicate the statistical significance using Student’s t-test: *, 0.01 < P-value < 0.05; **, 0.001 < P-value < 0.01. **(D)** Subcellular localization of SlERF-D7 in the nucleus of *Nicotiana benthamiana* epidermal cells.

### *SlERF.D7* is a nuclear-localized gene that responds positively to exogenous auxin and ethylene treatment

Next, we studied the subcellular localization of SlERF.D7 by transiently expressing *SlERF.D7::GFP* construct in *N. benthamiana* leaves. The *CaMV35S*-driven *SlERF.D7::GFP* fusion protein was found to be exclusively localized to the nucleus, consistent with its putative role in transcriptional regulation (Fig. 1D; Figure S1). To further characterize the SlERF.D7 protein, we determined its transactivation potential via transient expression assay using a GAL4-responsive reporter system. As hypothesized, SlERF.D7 displayed a strong transcriptional activation potential in the yeast system (Fig. 2A).

**Figure 2.**
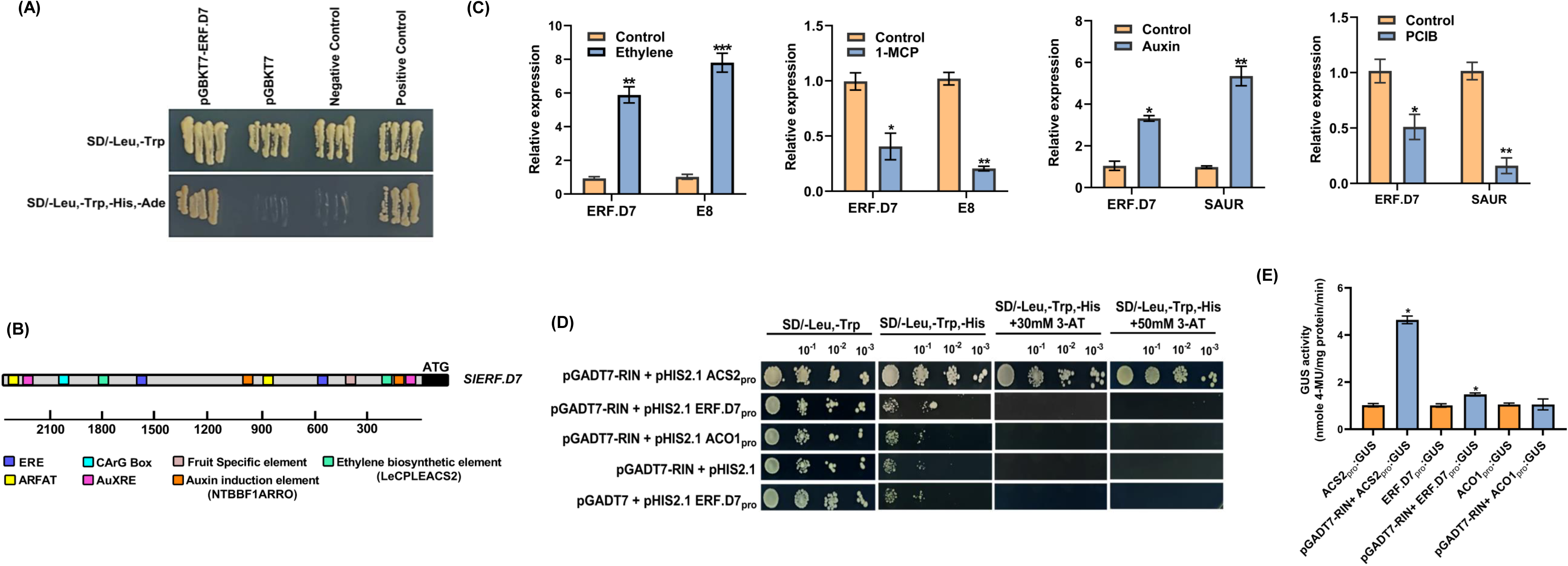
RIN-independent and ethylene- and auxin-dependent mode of action of SlERF.D7. **(A)** Analysis of transactivation potential of SlERF.D7 in yeast by growing transformants on synthetic dextrose (SD) media-lacking leucine (Leu), tryptophan (Trp), histidine (His), and Adenine (Ade). **(B)** The presence of CArG Box, a putative fruit-specific element in addition to putative ethylene and auxin response elements in the promoter of *SlERF.D7* gene. The *cis*-acting regulatory elements identified are represented by different color boxes. **(C)** qRT-PCR analysis of *SlERF.D7* transcripts in total RNA samples extracted from wild-type (WT) mature green fruit samples treated with 100 µM Ethrel, 100 µM 1-MCP, 100 μM IAA and 100 µM PCIB. Error bars, mean ± SD of three biological replicates. Asterisks indicate the statistical significance using Student’s t-test: *, 0.01 < P-value < 0.05; **, 0.001 < P-value < 0.01. E8, an ethylene response gene; SAUR68, an auxin response gene. **(D, E)** Identification of DNA binding activity of RIN to the promoter of *SlERF.D7* with (d) growth performance of transformants on SD/−Leu−/Trp/−His medium containing 50 mM 3-AT. Binding of RIN to the promoter of tomato *ACC synthase2* (*ACS2*) gene-: positive control, binding of RIN to the promoter of tomato *ACC oxidase1* (*ACO1*) gene-: negative control (d) *In vivo* interaction study of RIN to promoters of *SlERF.D7*, *SlACS2,* and *SlACO1* via GUS reporter assays in tobacco leaves.

We have previously observed the activation of ethylene-related ripening-associated genes by exogenous auxin treatment and *vice-versa* (Kumar et al., 2012a, b; SravanKumar et al., 2018). Mining of the 2.5-kb upstream sequence of *SlERF.D7* for *cis*-acting regulatory elements using the PLACE/signal search tool (http://www.dna.affrc.go.jp/PLACE/signalscan.html) revealed the presence of two putative Ethylene Response Elements (ERE-CCGAC) and two Auxin Response Elements (AuxRE-TGTCTC) in its promoter region (Fig. 2B). Furthermore, a putative fruit-specific element variant with a sequence motif TCTTCACA was also identified in the promoter region. We next investigated the influence of these two hormones on the transcriptional regulation of *SlERF.D7.* Exogenous treatment of both ethylene and auxin led to induction of *SlERF.D7* mRNA levels (Fig. 2C). Furthermore, a decline in its mRNA levels upon treatment with either 1-MCP (100 µM), an inhibitor of ethylene perception, or PCIB (100 µM), an antagonist of auxin function, validated the positive effect of both ethylene and auxin on *SlERF.D7* transcription (Fig. 2C). We estimated the efficacy of hormonal treatments by evaluating the transcript abundance of known ethylene (*E8*) and auxin (*SAUR*) responsive genes.

### SlERF.D7 is not transcriptionally activated by RIN

MADS-box proteins such as RIN, TAGL1, FUL1, and FUL2 have been previously reported to bind to CArG box elements in the promoters of ripening-related genes such as *SlACS2* and *SlPSY1* (Fujisawa *et al*., 2011; Li *et al*., 2019). The presence of a CArG box element in the upstream region of *SlERF.D7* (Fig. 2B) instigated us to examine its direct regulation by the ripening master regulator RIN using yeast-1-hybrid (Y1H) assay. Failure of survival of yeast cells containing both RIN protein and *SlERF.D7* promoter on selection media suggested no direct binding of RIN to the promoter. In contrast, yeast cells with RIN-activated *SlACS2* promoter, the positive control taken in the study, survived on the selection media, thereby conferring resistance in the cells to successfully grow on the selection medium (Fig. 2D). To further corroborate this result in vivo, we conducted a transient transactivation assay of *SlERF.D7* promoter by RIN in *N. benthamiana* leaves using GUS and GFP reporter systems. For this purpose, four-week-old Nicotiana leaves were co-infiltrated with the effector construct carrying the *RIN* coding sequence driven by the *CAMV35S* promoter. The reporter constructs were GUS or GFP coding sequences driven by *SlERF.D7* promoter. As noticed in the Y1H assay, there was no significant alteration in *SlERF.D7*-driven GUS activity or GFP fluorescence, in contrast to the positive control *SlACS2* driven enhanced GUS activity and GFP fluorescence transactivation assays confirming that RIN is incapable of binding to the *SlERF.D7* promoter (Fig. 2E). These observations indicated that *SlERF.D7* induction during fruit ripening is independent of RIN.

### Ripening-specific over-expression and silencing of *SlERF.D7* display dramatic but contrasting ripening-related changes in the transgenic fruits

To further elucidate the role of SlERF.D7 in tomato fruit ripening, we generated *SlERF.D7* over-expression and RNAi knock-down transgenic lines in tomato cultivar ‘Pusa Ruby’ under a ripening-specific *RIP1* promoter, characterized in our lab earlier (Fig. 3A) (Agarwal *et al*., 2017). *RIP1* gene is activated at post breaker stage of ripening. A total of 12 independent over-expression and eight independent silencing transgenic lines were obtained (Table. S2). Based on fruits exhibiting the highest and the lowest transcript levels of *SlERF.D7,* we selected two homozygous T_2_ generation representative transgenic lines for overexpression (OE) and silencing (RNAi) for further characterization. At the molecular level, the selected two OE lines, *SlERF.D7-OE#3* and *SlERF.D7-OE#11* exhibited the significantly elevated transcript accumulation at Br+3 and Br+10 stages compared to the tissue culture grown wild-type control fruits (Figure 3d). Similarly, two RNAi lines, *SlERF.D7-RNAi#1* and *SlERF.D7-RNAi#4,* displayed the strongest suppression of its transcripts, showing only 15-20% accumulation of its wild-type fruits transcripts at the Br+10 stage (Fig. 3B). No visible phenotypic differences in plant height, flower phenotype, fruit morphology, fruit set, and fruit development in both OE and RNAi lines compared to the wild-type control plants were observed by us. Considering the ripening-related expression pattern of *SlERF.D7*, we next examined the phenotype of OE and RNAi transgenic fruits in detail. *SlERF.D7*-*OE#3* and *SlERF.D7-OE#11* OE genotypes developed an intense red color at the Br+10 stage compared to the wild-type fruits, whereas *SlERF.D7-RNAi#1* and *SlERF.D7-RNAi#4* fruits exhibited inhibited red color development and failed to turn fully red. The RNAi fruits developed a mottled ripening pattern with partial degradation of chlorophyll at the final ripe stage (Fig. 3C). However, we noticed discernable phenotypic alterations pertaining to fruit softening in the fruits of *SlERF.D7 OE* lines as both *SlERF.D7*-*OE#3* and *SlERF.D7-OE#11* fruits displayed wrinkling of the outer pericarp tissue, possibly accounting for early signs of cell wall loosening than their wild-type controls (Fig. 3C). Off-wine analysis of *SlERF.D7* OE and RNAi fruits exhibited the same ripening pattern, with OE lines fruits displaying signs of premature ripening compared to the wild-type control fruits. Contrastingly, *SlERF.D7* RNAi silenced fruits showed less pigment accumulation coupled with increased firmness than their same-age wild-type control fruits (Fig. 3D). Altogether, we observed contrasting fruit phenotypes regarding fruit color and chlorophyll degradation in OE and RNAi lines. These observations further indicated that the degradation of chlorophyll, synchronous with the synthesis of carotenoids and cell wall degradation during the ripening process, is directly targeted by *SlERF.D7*.

**Figure 3.**
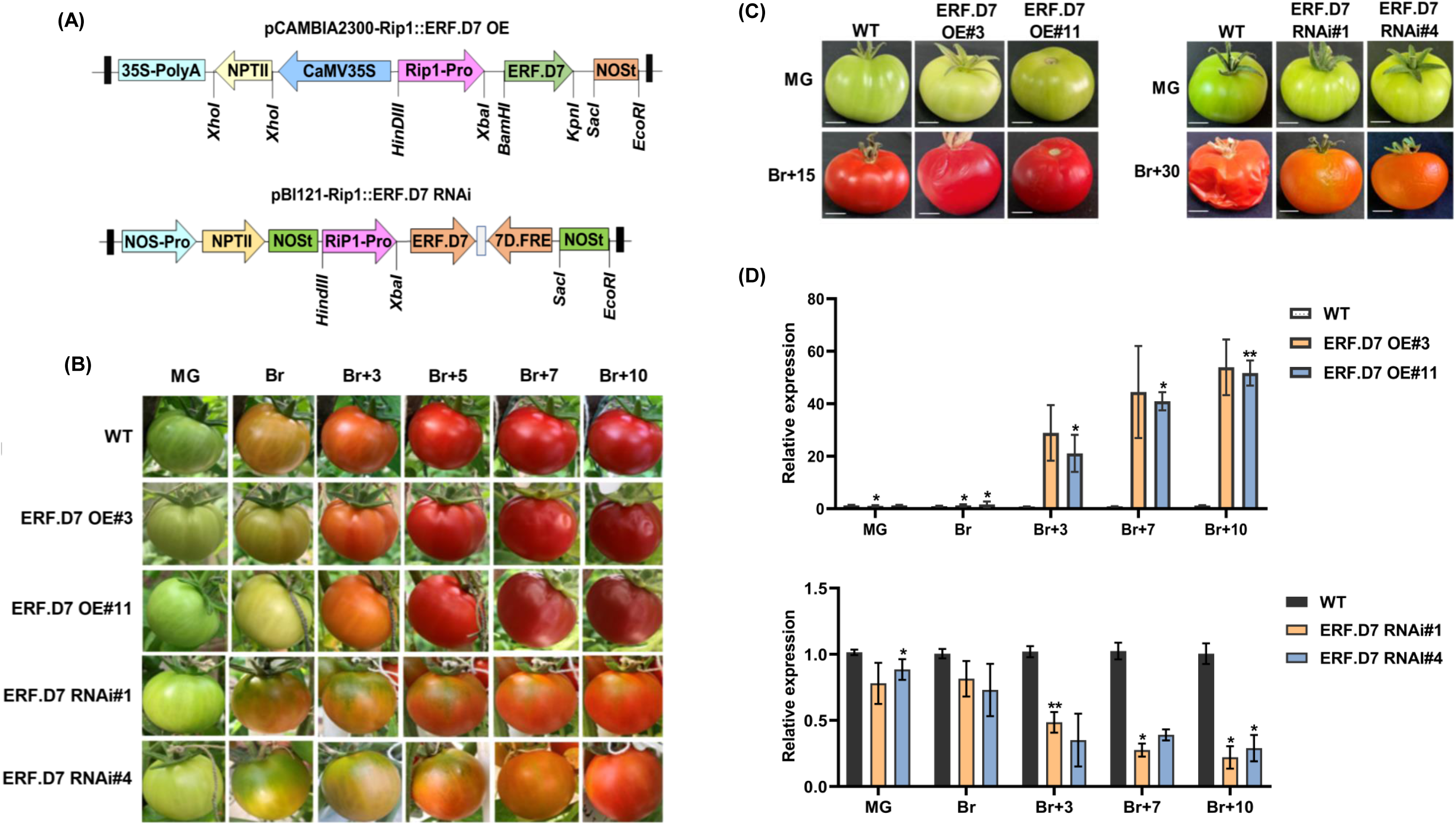
Morphological and molecular characterization of *SlERF.D7* overexpression and RNAi lines. **(A)** Vector diagram of *pCAMBIA2300-Rip1::SlERF.D7* OE (over-expression) and *pBI121-Rip1::SlERF.D7* RNAi (silencing) construct. **(B)** Phenotypic analysis of wild-type (WT), *SlERF.D7* OE, and RNAi line fruits at mature green (MG), breaker (Br), 3-day after Br (Br+3), 5-day after Br (Br+5), and 7-day after Br (Br+7). **(C)** Off wine phenotypic assessment of wild-type (WT), *SlERF.D7* OE and RNAi line fruits harvested at the mature green (MG) stage and photographed till the first sign of shriveling appears, 15-day post Br (Br+15) in *SlERF.D7* OE lines and 30-day post Br (Br+30) in *SlERF.D7* RNAi line. **(D)** Transcript levels of *SlERF.D7* in wild-type (WT), OE and RNAi transgenic fruit analysed at mature green (MG), breaker (Br), 3-day after Br (Br+3), 7-day after Br (Br+7), and 10-day after Br (Br+10) by qRT-PCR with *Actin* gene as an internal control. Error bars mean ±SD of three biological replicates. Asterisks indicate the statistical significance using Student’s t-test: *, 0.01 < P-value < 0.05; **, 0.001 < P-value < 0.01.

### Down-regulation of *SlERF.D7* leads to reduced pigment accumulation and enhanced fruit firmness

To investigate the possible cause of altered pigmentation in *SlERF.D7* OE and RNAi fruits, we performed HPLC-based carotenoid profiling in wild-type and transgenic line pericarp tissue at MG, Br+3 Br+7, and Br+10 stages. In terms of lycopene accumulation, a 55-60% reduction was observed in SlERF.D7-RNAi#1 and SlERF.D7-RNAi#4 fruits compared to wild-type fruits at the Br+10 stage. In contrast, the fruits of *SlERF.D7-OE#3* and *SlERF.D7-OE#11* lines displayed a 40-45% enhancement in lycopene levels at the Br+10 stage compared to wild-type fruits, consequently imparting them an intense red color phenotype (Fig. 4A). Concomitantly, a sharp increase in β-carotene content was detected in *SlERF.D7* RNAi fruits at the Br+10 stage, keeping with an orange fruit phenotype (Fig. 4B). Similar to an opposite trend observed for lycopene content in OE and RNAi lines, β-carotene content also exhibited contrasting profiles in the fruits of the two sets of transgenic plants. To uncover the molecular basis of this modulation in carotenoid composition, we analyzed the transcript level of genes involved in the carotenoid biosynthetic pathway of pericarp tissue at different stages during fruit ripening by qRT-PCR. Transcript abundance of *phytoene synthase*, *PSY1*, a key regulator of the carotenoid pathway, was severely repressed in *SlERF.D7-RNAi#1* and *SlERF.D7-RNAi#4* fruits at B+3 and later ripening stages, concomitant to the silencing pattern of this gene in the RNAi lines (Fig. 4C). A similar reduction in mRNA levels of *phytoene desaturase* (*SlPDS)*, was also observed in *SlERF.D7* RNAi fruits (Fig. 4C). In contrast, transcripts accumulation of all the three lycopene-β-cyclases (β-*LYC1*, β-*LYC2,* and *CYC*-β) were significantly upregulated in *SlERF.D7* RNAi fruits as compared to their wild-type counterparts, probably responsible for the elevated β-carotene accumulation in these transgenic fruits (Fig. 4C). Likewise, a substantial increase in the transcript levels of *SlPSY1* and *SlPDS* with a coordinated decline in expression levels of *β-LYC1*, *β-LYC2,* and *CYC*-β in *SlERF.D7-OE#3* and *SlERF.D7-OE#11* fruits, at Br+3 and later stages, accounted for the enhanced carotenoid accumulation in these fruits (Fig.4C). The data indicate that repression of *SlERF.D7* culminates in modified lycopene to β-carotene ratio via the alteration in expression levels of key carotenoid pathways genes such as *PSY1*, *PDS,* and lycopene β-cyclases.

**Figure 4.**
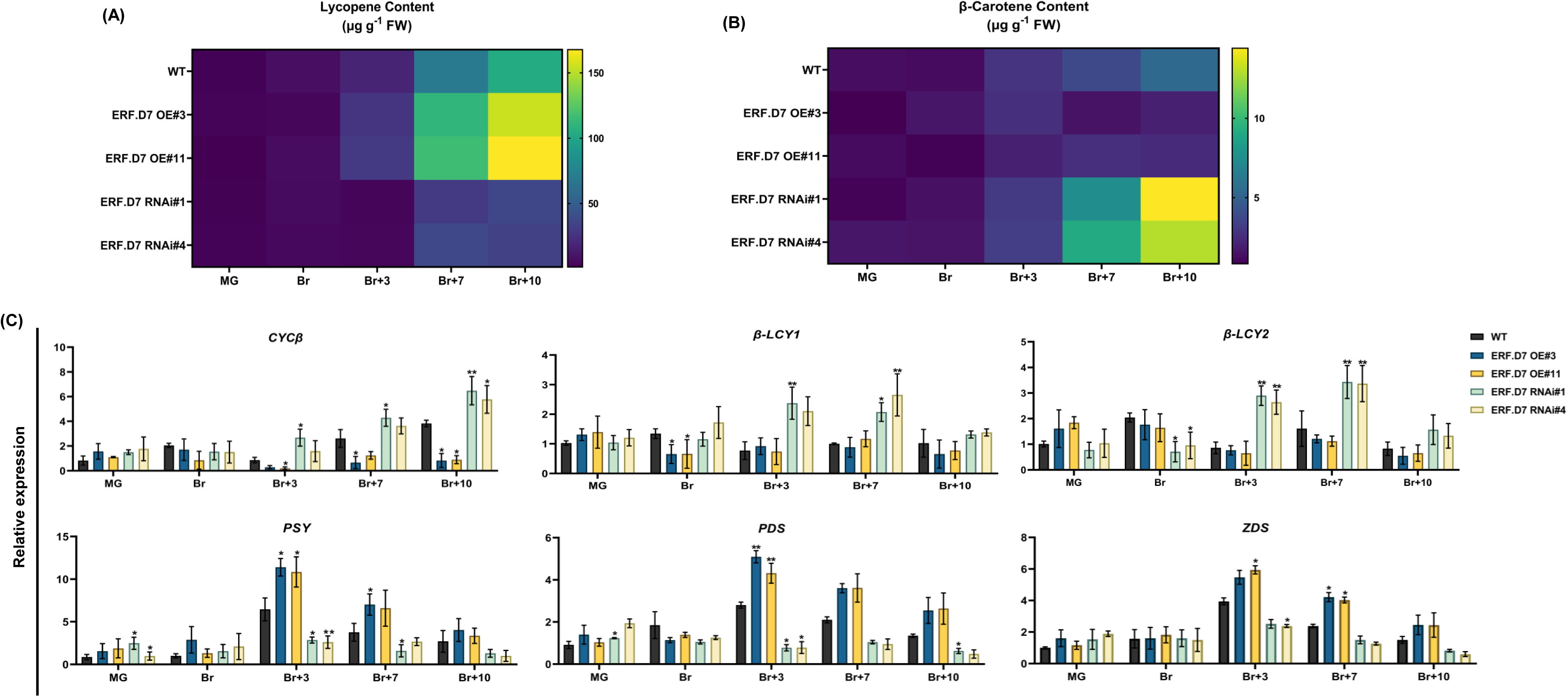
Pigment accumulation assessment in *SlERF.D7* OE and RNAi fruits. **(A,B)** Estimation of **(A)** lycopene and **(B)** β-carotene in wild-type (WT) and *SlERF.D7* OE and RNAi lines at different stages of fruit ripening. **(C)** qRT-PCR analysis of carotenoid biosynthetic pathway genes in wild-type (WT) and *SlERF.D7* OE and RNAi tomato lines at mature green (MG), breaker (Br), 3-day after Br (Br+3), 7-day after Br (Br+7) and 10-day after Br (Br+10) with *Actin* gene as an internal control. Error bar means ±SD of three biological replicates. Asterisks indicate the statistical significance using Student’s t-test: *, 0.01 < P-value < 0.05; **, 0.001 < P-value < 0.01. β-LCY1, β-LCY2, CYC-β lycopene b-cyclases; PSY1 phytoene synthase; PDS phytoene desaturase; ZDS, carotenoid desaturase.

Given that fruit softening is another critical parameter of fruit ripening, we assessed the progression of firmness in *SlERF.D7* OE and RNAi lines fruits from MG to Br+10 stage. Acceleration of 35% in fruit softening was detected in *SlERF.D7-OE#3* and *SlERF.D7-OE#11* fruits at the Br+3 stage compared to their wild-type control (Fig. 5A). On the other hand, *SlERF.D7* RNAi silenced fruits were associated with a noticeable increase in fruit firmness, which reached a maximum of two to three times higher than that in wild-type fruit at the Br+7 stage (Fig. 5A). To further substantiate these findings at the genetic level, we examined the mRNA abundance of cell wall modifying enzymes such as polygalacturonase-2a (*PG2A*) and pectate lyase (*PL*) using qRT-PCR. *SlERF.D7-OE#3* and *SlERF.D7-OE#11* lines fruits accumulated higher *PG2A* and *PL* transcripts than the wild-type fruits at the Br+3 and Br+7 stages, consistent with the shriveled appearance of these fruits (Fig. 5B). Contrastingly, transcripts levels of these two genes were significantly inhibited at the B+3 stage onwards in the fruits of *SlERF.D7* RNAi lines compared to their wild-type controls. Altogether, data indicate that once the ripening program initiates, it proceeds at a much more intense rate in *SlERF.D7* OE transgenic fruits and milder in the *SlERF.D7* RNAi fruits in comparison to their wild-type control counterparts.

**Figure 5.**
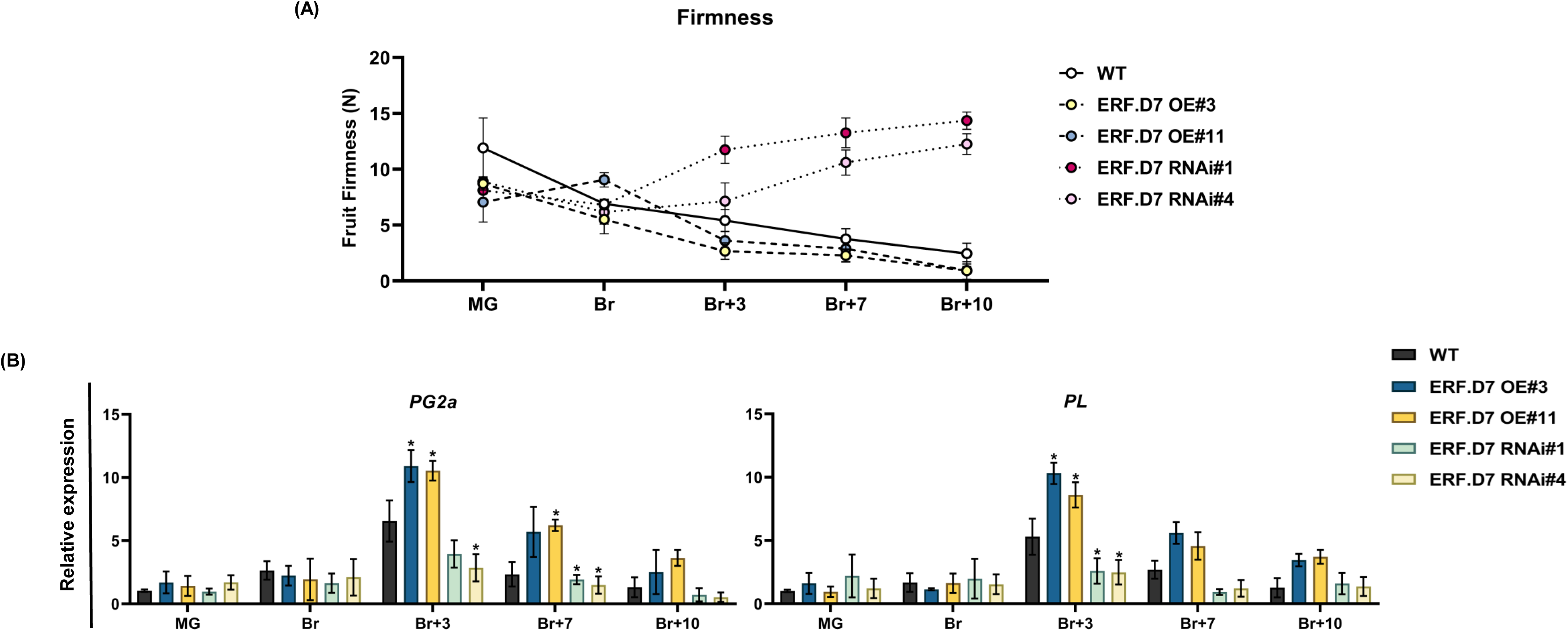
Modulations in levels of fruit firmness in *SlERF.D7* OE and RNAi fruits. **(A)** Fruit firmness analysis in WT, *SlERF.D7* OE, and RNAi line fruits at different stages of ripening, with fruits being harvested at mature green (MG) stage. A total of 15 fruits was used for each measurement, and the values shown are the means ±SD. **(B)** qRT-PCR expression of polygalacturonase gene (*SlPG2A*) and pectate lyase (*SlPL*) at mature green (MG), breaker (Br), 3-day after Br (Br+3), 7-day after Br (Br+7), and 10-day after Br (Br+10) in *SlERF.D7* OE, RNAi and WT fruits with *GAPDH* and *Actin* were used as the internal controls. The Error bars represent ±SD of three biological replicates. Asterisks indicate the statistical significance using Student’s t-test: *, 0.01 < P-value < 0.05; **, 0.001 < P-value < 0.01.

### Ethylene emission and perception are altered in *SlERF.D7* OE fruits

Since increased ethylene output is instrumental in determining the speed of ripening, carotenoid accumulation, and fruit firmness, we next assessed ethylene production in *SlERF.D7* OE and RNAi fruits from MG to B+10 stage. A 40-45 percent rise in ethylene emission was recorded in *SlERF.D7-OE#3* and *SlERF.D7-OE#11* transgenic fruits compared to the same-stage wild-type control fruits. In contrast, significantly lower ethylene production was observed in *SlERF.D7* RNAi silenced fruits at the B+3 stage. The atypical ethylene maxima observed at the Br+3 stage in the wild-type fruits were missing in the RNAi lines fruits (Fig. 6A). Assessment of the expression of key ethylene biosynthetic genes by qRT-PCR revealed elevated *SlACO1*, *SlACS2,* and *SlACS4* transcripts in *SlERF.D7* OE fruits at B+3 or later ripening stages (Fig. 6C). On the other hand, *SlERF.D7* RNAi fruits displayed reduced mRNA levels at Br+3, Br+7, and Br+10 stages compared to their same-stage wild-type counterparts (Fig. 6C). Interestingly, to compensate for low ethylene production, exogenous ethylene application to SlERF.D7 RNAi fruits at the mature green (MG) stage failed to reverse the inhibited ripening phenotype (Fig. 6B). To corroborate this, we conducted an expression profiling of genes involved in the ethylene signalling pathway in wild-type, *SlERF.D7* RNAi, and OE fruits at different stages of ripening. We noticed a drastic reduction in the transcripts accumulation of Ethylene Receptors such as *ETR2*, *ETR3* (*NR*), *ETR4,* and *ETR5* in post breaker stages of *SlERF.D7* RNAi fruits as compared to the wild-type fruits (Fig. 6C). Among the other important ethylene signalling genes, *ethylene-insensitive 2* (*EIN2*) and *EIN2-like* (*EIL2*) were also down-regulated during ripening in *SlERF.D7* RNAi line fruits (Fig.6C). To further narrow down to ERFs, altered but mostly opposite expression patterns of numerous ERFs were observed in *SlERF.D7* OE and RNAi fruits to each other.

**Figure 6.**
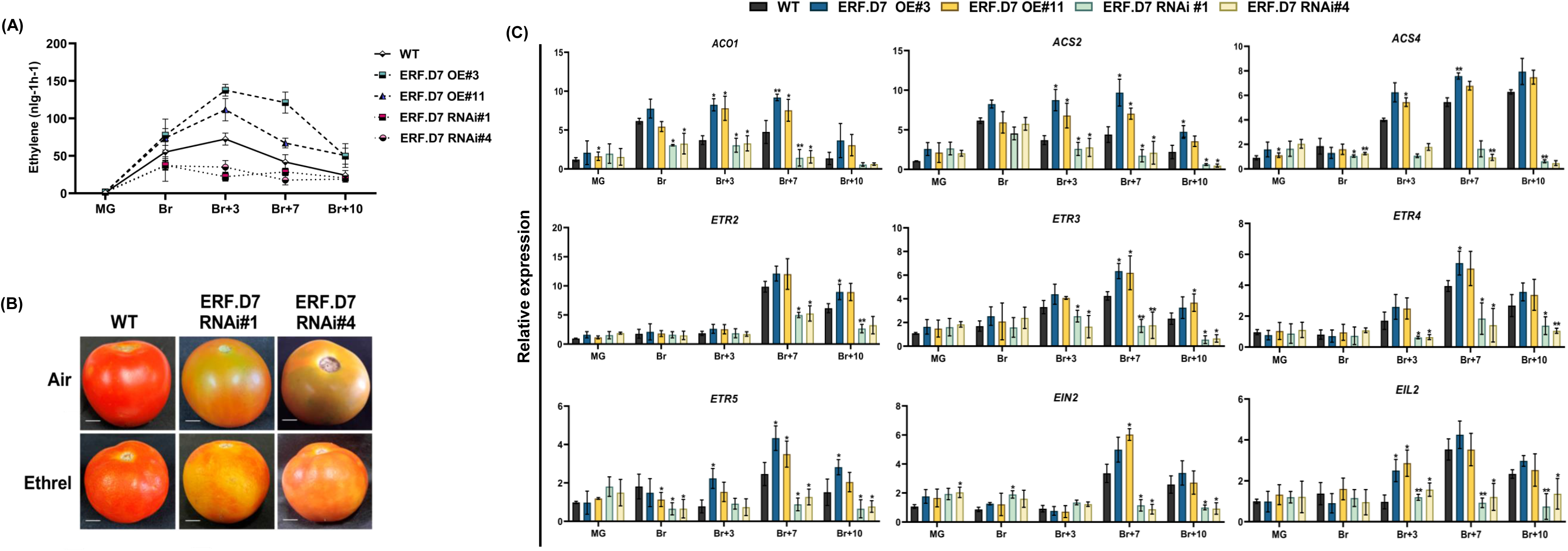
Alterations in ethylene biosynthesis and perception in *SlERF.D7* transgenic fruits. **(A)** Ethylene production of WT and *SlERF.D7* OE and RNAi fruits assessed at mature green (MG), breaker (Br), 3-day after Br (Br+3), 7-day after Br (Br+7), and 10-day after Br (Br+10) stages. Values represent the means of at least five individual fruits. **(B)** Exogenous ethylene treatment on wild-type (WT) and *SlERF.D7* RNAi fruit. Mature green fruits from WT and *SlERF.D7* RNAi lines were injected a buffer solution containing 10 mM MES, pH 5.6, sorbitol (3% w/v) and 100 µM Ethrel (2-Chloroethylphosphonic Acid, 40% Solution, SRL Diagnostics). After the treatment, fruits were incubated in a culture room at 26°C, under a 16 h light/8 h dark cycle with a light intensity of 100 μmol m^-2^ s^-1^ and photographed for 7 days. **(C)** qRT-PCR expression of ethylene biosynthesis and perception pathway genes at mature green (MG), breaker (Br), 3-day after Br (Br+3), 7-day after Br (Br+7), and 10-day after Br (Br+10) in *SlERF.D7* OE, *SlERF.D7* RNAi and WT fruits with *Actin* gene as an internal control. Error bars represent ±SD of three biological replicates. Asterisks indicate the statistical significance using Student’s t-test: *, 0.01 < P-value < 0.05; **, 0.001 < P-value < 0.01. ACO1, aminocyclopropane-1-carboxylic acid oxidase 1; ACS2 and ACS4 aminocyclopropane-1-carboxylic acid synthases; ETR2, ETR3, ETR4, and ETR5 ethylene receptors; EIN2 ethylene signalling protein; EIL2 EIN2-like protein.

Five ERF genes (*ERF.A3, ERF.C1, ERF.E1, ERF.E2,* and *ERF.E4*) displayed up-regulation at the onset of ripening starting from the Br stage in *SlERF.D7* OE lines (Fig. S3). In contrast, a concomitant decrease in transcripts of these five ERFs was observed in *SlERF.D7* RNAi fruits at post-breaker stages. By contrast, the mRNA abundance of three ERFs (*SlERF.B1, SlERF.B2,* and *SlERF.B3*) exhibited enhanced mRNA abundance in *SlERF.D7* RNAi line fruits (Fig. S3). To rule out any possibility of off-target effects *SlERF.D7* transgenic line, we further investigated the expression profiles of all SlERF.D clade members. We found no significant changes in any SlERF.D clade member’s transcript abundance in transgenic fruits (Fig.S4). In conclusion, these results signify that ethylene biosynthesis coupled with ethylene perception and signalling contributes to the altered ripening phenotypes observed in the OE and RNAi lines.

### Transcription of key ripening regulators is modulated in *SlERF.D7* transgenic line fruits

To unveil the possible molecular mechanism responsible for this altered ripening phenotypes of *SlERF.D7* transgenic lines, we assessed the transcript profiles of major ripening regulator genes at different stages of ripening. Compared to wild-type fruits, *RIN* and *CNR* mRNA levels were significantly reduced at Br+7 and Br+10 stage in the RNAi fruits (Fig.7). Likewise, 50-60% reduction in the transcripts of *FRUITFUL1* (*FUL1)* and *FRUITFUL2* (*FUL2)* were observed in *SlERF.D7* RNAi fruits at post breaker stages (Fig. 7). Similar inhibition was also observed in mRNA levels of *NOR, SlAP2a,* and *TAGL1* genes in the RNAi lines fruits (Fig. 7). The altered expression pattern of most of these genes in *SlERF.D7* transgenic fruits is consistent with the delayed onset of ripening observed in the RNAi lines. Correspondingly, *SlERF.D7* OE fruits accumulated elevated transcripts of *RIN*, *CNR*, *NOR, AP2A,* and *TAGL1*, compared to the wild-type control fruits, at post breaker stages.

**Figure 7.**
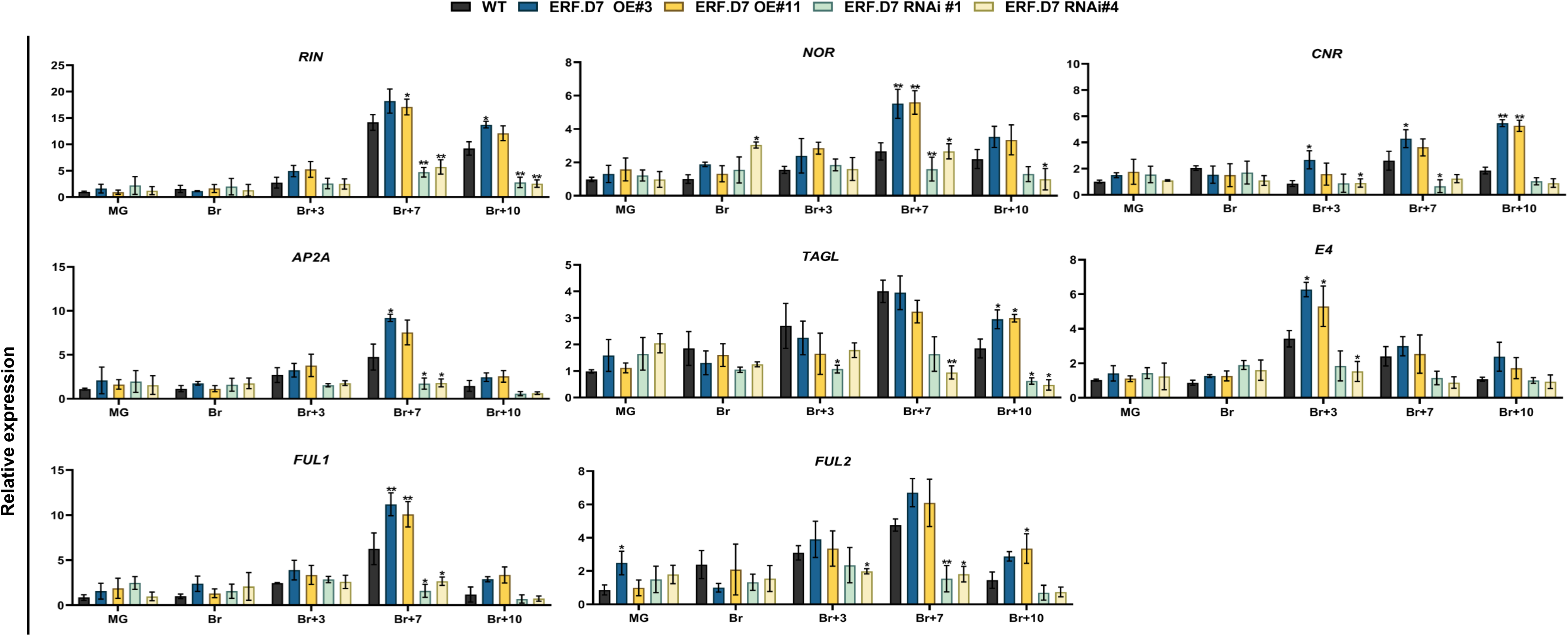
Transcript profiling of key ripening regulator genes in wild-type (WT) and *SlERF.D7* OE and RNAi tomato lines during fruit ripening. Total RNA was extracted from mature green (MG), breaker (Br), 3-day after Br (Br+3), 7-day after Br (Br+7), and 10-day after Br (Br+10) stages of *SlERF.D7* OE, SlERF.D7 RNAi and WT fruits. The relative mRNA levels of each gene was normalized using the *Actin* gene as an internal control. Error bars represent ±SD of three biological replicates. Asterisks indicate the statistical significance using Student’s t-test: *, 0.01 < P-value < 0.05; **, 0.001 < P-value < 0.01. RIN ripening inhibitor; NOR non-ripening; CNR colorless non-ripening; AP2a APETALA2/ERF gene; TAGL1 tomato AGAMOUS-LIKE 1; E4 ethylene-responsive and ripening regulated genes; FUL1 and FUL2 Fruitful MADS-box transcription factor homologs.

### SlERF.D7 alters fruit ripening by influencing auxin sensitivity

Because *SlERF.D7* positively responded to IAA treatment, and the fact that *SlERF.D7* RNAi fruits closely mimicked the ripening phenotype of *SlARF2A* and *SlARF2B* RNAi (or OE) fruits reported earlier (Hao *et al*., 2015; Breitel *et al*., 2016*a*), we next assessed the levels of IAA in transgenic fruits at MG, Br and Br+7 stages. Remarkably, a significant increase in IAA concentration was observed in *SlERF.D7* RNAi fruits at the Br+7 stage (Fig. 8A). In contrast, the OE lines showed slightly decreased IAA levels. Considering the modulated IAA levels in transgenic fruits and the known roles of tomato ARFs in ripening, we then examined the expression profile of all tomato ARF gene family members by qRT-PCR (Kumar *et al*., 2015) (Fig. S5). Strikingly, the transcript accumulation of *SlARF2* orthologs was significantly affected in the transgenic fruits. Transcription of no other ARF gene family members was affected in *SlERF.D7* transgenic fruits (Fig. S5). In particular, *SlARF2A* displayed a dramatic down-regulation in *SlERF.D7-RNAi#1* and *SlERF.D7-RNAi#4* fruits at the Br+3 stage, whereas *SlARF2B* exhibited a slight reduction in its mRNA levels at this stage (Fig. 8B). However, transcript accumulation of *SlARF2B* was reduced to 30% in *SlERF.D7* RNAi fruits at the Br+7 stage (Fig. 8A). On a similar level, *SlERF.D7* OE line fruits showed a significant upregulation in the expression of *SlARF2A* at post breaker stages. These results signified a possible role of SlERF.D7 in controlling ripening traits by moderating auxin responses, plausibly through SlARF2 orthologs, in tomato fruits.

**Figure 8.**
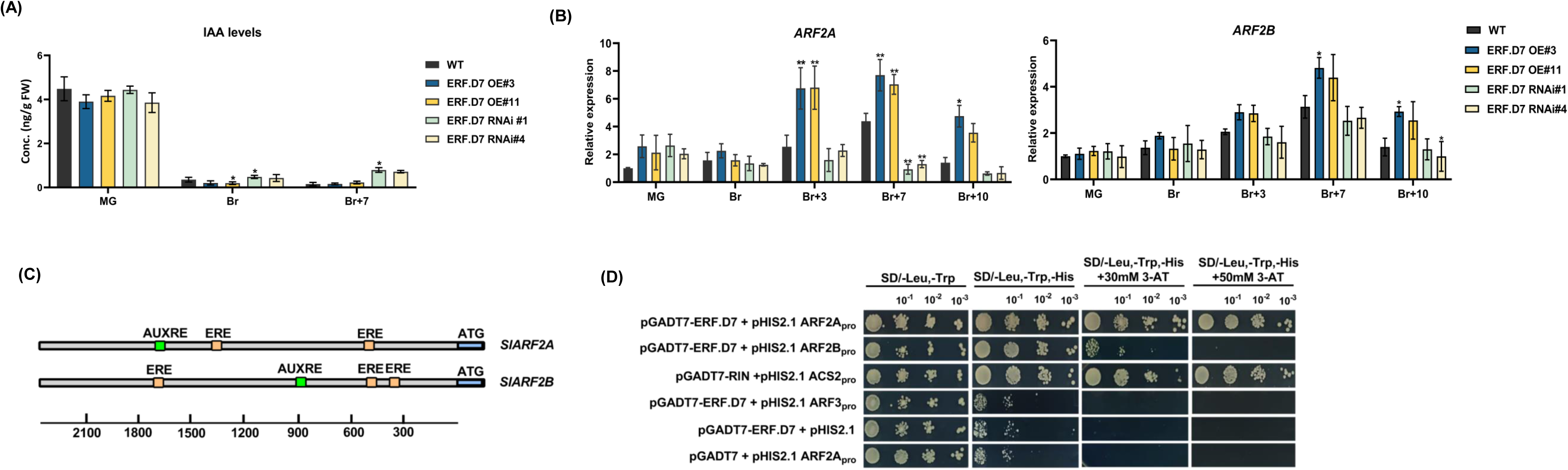
Altered auxin levels and ARF2 orthologs transcription in *SlERF.D7* transgenic fruits. **(A)** Determination of IAA levels during fruit ripening in wild-type (WT), *SlERF.D7* OE, and RNAi line fruits at MG, breaker (Br), and 7-day post breaker (Br + 10) stages. **(B)** Expression profiling of *SlARF2* orthologs, *SlARF2A* and *SlARF2B* in transgenic fruits of *SlERF.D7* OE and RNAi lines during different fruit maturation stages, mature green (MG), breaker (Br), 3-day after Br (Br+3), 7-day after Br (Br+7) and 10-day after Br (Br+10). The relative mRNA levels of both the genes were normalized using *Actin* and *GAPDH* genes as an internal control. Error bars represent ±SD of three biological replicates. Asterisks indicate the statistical significance using Student’s t-test: *, 0.01 < P-value < 0.05; **, 0.001 < P-value < 0.01. **(C)** Analysis of putative ethylene and auxin-responsive cis-elements in the 2.5-kb promoter region of *SlARF2A* and *SlARF2B* genes. **(D)** Yeast-1-hybrid (Y1H) analysis of binding of SlERF.D7 protein to putative ethylene-responsive elements in the promoter regions of *SlARF2A* and *SlARF2B* genes displaying growth performance of transformants on SD/−Leu−/Trp/−His medium containing 50 mM 3-AT. Binding of RIN to the promoter of *SlACS2* gene-: positive control, binding of *SlERF.D7* to the promoter of *SlARF3* gene-: negative control.

### SlERF.D7 binds to and activates the promoters of *SlARF2A/B*

Because of the observed alterations in the transcript level of *SlARF2A* and *SlARF2B* in *SlERF.D7* transgenic lines fruits, we hypothesized that SlERF.D7 might bind to the promoters of the two *ARF2* orthologs. *In silico* analysis of 2.5-kb promoter sequences of *SlARF2A* and *SlARF2B* revealed the presence of two and three conserved Ethylene Responsive Elements (ERE), respectively (Fig. 8C). These elements are the putative targets of ERF-type transcription factors in plants. To validate this assumption, we performed a yeast one-hybrid assay. A *pGAD::SlERF.D7* plasmid (containing the SlERF.D7 putative DNA domain fused to the GAL4 active domain) and *pHIS*-cis-acting reporter construct (2-kb PCR amplified promoter regions of *SlARF2A* and *SlARF2B*) were co-transformed into yeast strain Y187. The results displayed a strong binding of SlERF.D7 to the *SlARF2A* promoter, whereas a weak interaction between this protein and *SlARF2B* promoter was noticed (Fig. 8D). As indicated by the modulations in the transcript levels of these reporter genes, *SlERF.D7* could directly bind to the promoters of *SlARF2A* and *SlARF2B* and regulate the expression of these target genes.

To further determine SlERF.D7-mediated direct activation of *SlARF2* genes *in-planta*, we evaluated GUS and GFP activity driven by *SlARF2A* and *SlARF2B* promoter fusion reporter constructs in *N. benthamiana* leaves that also transiently expressed the *SlERF.D7* gene under the control of *CaMV35S* promoter (effector construct) (Fig. 9A,B). Transactivation of the *SlARF2A* promoter by SlERF.D7 significantly enhanced the GUS reporter activity. However, the transient co-expression of *SlARF2B* promoter and SlERF.D7 displayed less GUS activity elevation than *SlARF2A* and the positive controls (Fig. 9C). Similar observations were made with GFP fluorescence driven by transient co-expression of *SlARF2A* and *SlARF2B* reporter constructs and SlERF.D7 effector construct in tobacco leaves (Fig. 9B,D). The data indicate that SlERF.D7 regulates the expression of *SlARF2* orthologs by directly binding, most likely to the typical ERE elements, in their promoter regions.

**Figure 9.**
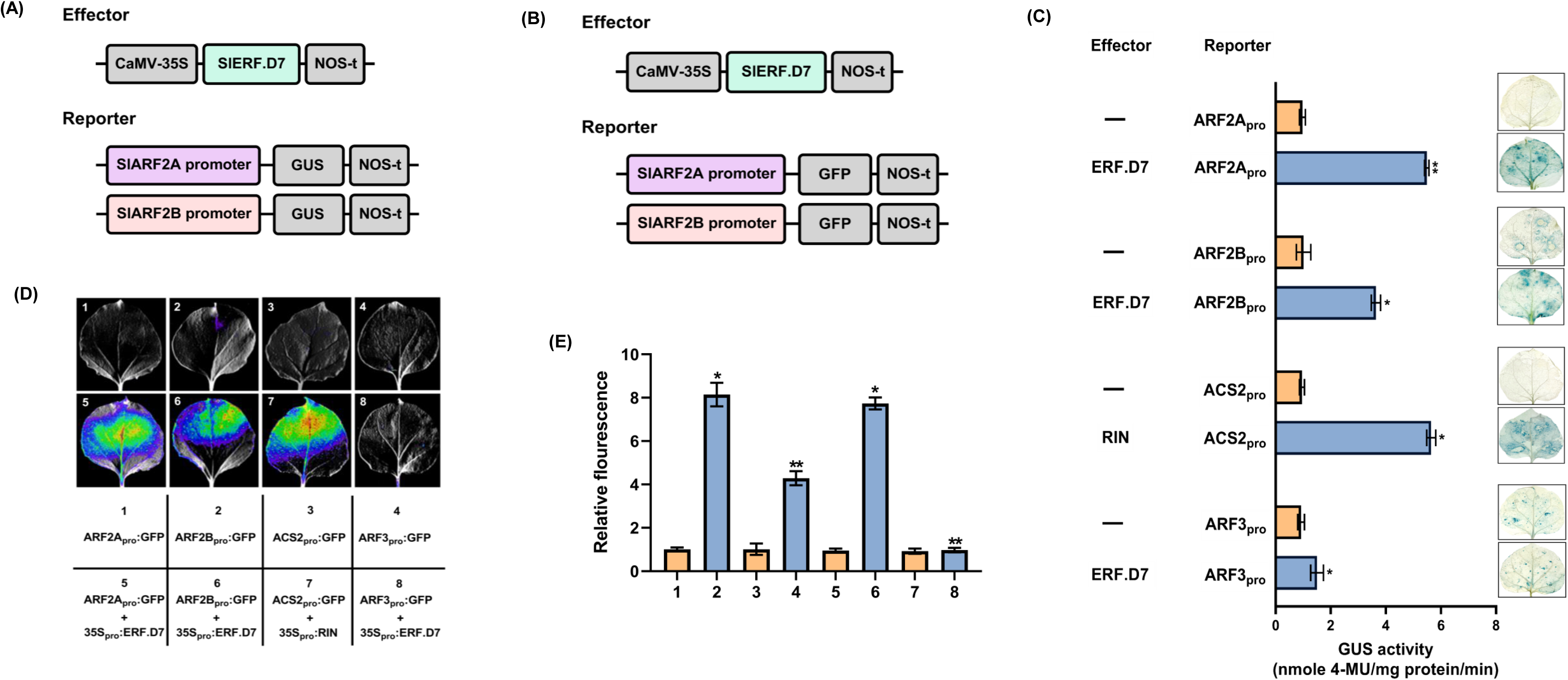
In vivo transactivation of SlARF2 promoters by SlERF.D7. **(A,B)** Schematic diagrams of the reporter and effector constructs. The GUS and GFP reporter plasmids contain the promoters of the *SlARF2A* and *SlARF2B*. The effector plasmids contain the *SlERF.D7* coding sequence under the control of the *CaMV35S* promoter. **(C)** Confirmation of the activation of SlARF2A and SlARF2B promoters by SlERF.D7 via the GUS complementation assay in *N. benthamiana* leaves. GUS assay was performed after 48 hr of infiltration. GUS activity is expressed in nmole 4-methylumelliferone mg^-1^ protein min^-1^. Activation of *SlACS2* promoter by RIN-: positive control, Activation of *SlARF3* promoter by SlERF.D7-: negative control. Error bars represent ±SD of three biological replicates. Asterisks indicate the statistical significance using Student’s t-test: *, 0.01 < P-value < 0.05; **, 0.001 < P-value < 0.01.(**D,E**) Transient expression assay shows that SlERF.D7 activates the expression of the GFP reporter gene. The bottom panel indicates the combination of reporter and effector plasmids infiltrated. The GFP reporter gene is driven by *ARF2A* and *ARF2B* promoters. Representative images of *N. benthamiana* leaves 48 h after infiltration captured via NightSHADE Plant Imaging System are shown. Activation of *SlACS2* promoter by RIN-: positive control, Activation of *SlARF3* promoter by SlERF.D7-: negative control. **(E)** Quantitative analysis of fluorescence intensity in three independent determinations was assessed. Error bars represent ±SD of three biological replicates. Asterisks indicate the statistical significance using Student’s t-test: *, 0.01 < P-value < 0.05; **, 0.001 < P-value < 0.01.

Although the transient GUS and GFP transactivation assays suggested a direct regulation of *SlARF2* promoters by SlERF.D7 protein, we further conducted an electrophoretic mobility shift assay (EMSA) to assess the ability of SlERF.D7 to directly bind to *SlARF2A* and *SlARF2B* promoters. Indeed SlERF.D7 exhibited direct binding to the DNA probe containing the ERE element present in the SlARF2A promoter, whereas the unlabeled promoter fragment displaced the binding of the labeled probe in a dose-dependent manner (Fig 10A). However, SlERF.D7 displayed weak binding to ERE motif in the SlARF2B promoter (Fig 10A). These results revealed the ability of SlERF.D7 to specifically bind to ERE motif in the *SlARF2A* promoter. Combined together, the data confirms that *SlARF2* promoters are direct targets of SlERF.D7 in planta.

**Figure 10.**
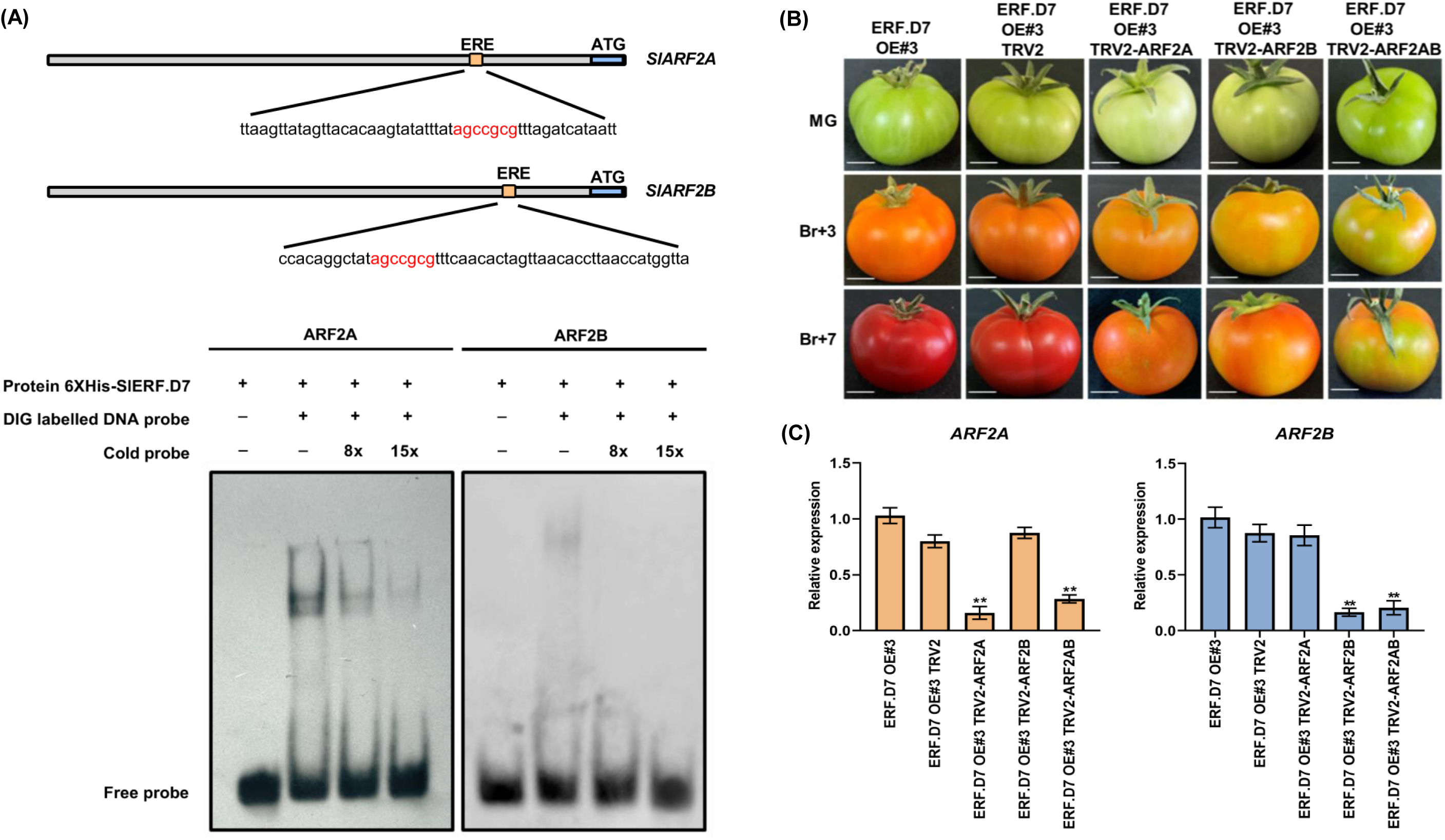
*SlARF2* promoters are a direct target of SlERF.D7. (A)) SlERF.D7 binding to the promoters of *SlARF2* harbouring ERE element. The WT probe containing the ERE motif was digoxigenin-labelled. Competition for SlERF.D7 binding was performed with 8x and 15x cold probes. The symbols - and + represent the absence or presence of the probes and 6X Histidine-tagged SlERF.D7 protein. **(B)** Phenotypes of *SlARF2A*, *SlARF2B* and *SlARF2AB* silenced fruits in *SlERF.D7* OE background. Un-infiltrated and vector-only (pTRV) fruits were used as the control. Photographs were taken at mature green (MG), 3-day post Br (Br+3), and 7-day post Br (Br+7) stages. **(C)** The silencing efficiency of the *SlARF2* orthologs in *SlARF2A*, *SlARF2B,* and *SlARF2AB* infiltrated fruits. The relative mRNA levels of both the genes were normalized using *GAPDH* and *Actin* gene as an internal control. Error bars represent ±SD of three biological replicates. Asterisks indicate the statistical significance using Student’s t-test: *, 0.01 < P-value < 0.05; **, 0.001 < P-value < 0.01.

### Silencing of *SlARF2* orthologs in *SlERF.D7* OE lines fruits partially reverse the fastened ripening phenotype

To confirm whether the elevated transcript levels of *SlARF2* orthologs were responsible for the enhanced carotenoid synthesis in *SlERF.D7* OE line fruits, we performed VIGS assays of *SlARF2A*, *SlARF2B,* and double knock-down of both *SlARF2A/B* in *SlERF.D7*-OE#3 fruits at the MG stage. A spotted ripening pattern was observed in *SlARF2A* single knock-down VIGS fruit, with the fruits remaining orange at the final maturation stage (Fig. 10B). Similarly, *SlARF2B* single knock-down VIGS fruits exhibited more distinct variegation of yellow and orange color on the outer pericarp and never achieved the intense red color of *SlERF.D7* OE fruits (Fig. 10B). The simultaneous double knock-down *SlARF2A/B* VIGS fruits displayed more severe ripening defects. These fruits showed mottled green and orange sectors, separated by distinct borders, contrasting with the uniform red color observed in *SlERF.D7* OE and *SlERF.D7* OE *TRV* control fruits, suggesting a redundant function of *SlARF2A* and *SlARF2B* in ripening (Fig. 10B). Previously, RNAi-mediated stable transgenic silencing lines of *SlARF2A*, *SlARF2B* along with a double knock-down *SlARF2A/B* have already been investigated for fruit ripening alterations by two groups (Hao *et al*., 2015; Breitel *et al*., 2016*a*). The authors have described a similar delayed ripening phenotype in RNAi lines as we noticed with a VIGS construct in *SlERF.D7-OE#3* fruits. Also, *SlERF.D7* RNAi silenced fruits in this study yield a similar but milder phenotype, which can be attributed to using a weaker but more ripening-specific *RIP1* promoter, instead of a constitutive *CaMV35S*, in VIGS experiments. To verify that the phenotype obtained was associated with the silencing of *ARF2A* and *ARF2B* in single and double VIGS silenced fruits, we analyzed the expression of these genes at the molecular level. A 70-75% reduction in transcript levels of *SlARF2* genes was observed in single and double knock-down VIGS fruits compared to *SlERF.D7-OE#3* fruits (Fig. 10C). Additionally, the mRNA abundance of *SlARF2B* in *SlARF2B* single and double knock-down VIGS fruits decreased to about 15-20% of their control OE fruits (Fig. 10C).

HPLC-based assessment of color change in VIGS fruit for lycopene and β-carotene further emphasized the difference in carotenoid production in *SlERF.D7-OE#3* and its SlARF2 orthologs VIGS fruits. The lycopene levels were significantly compromised in all the VIGS fruits, with Sl*ARF2A/B* double knock-down fruits displaying the maximum reduction (Fig. 11). In the context of β-carotene, an increase of 40-45% was observed in *SlARF2B* lines compared to *SlERF.D7-OE#3* line control fruits (Fig.11). Therefore, it can be interpreted that this decreased lycopene and increased β-carotene levels were responsible for the yellow-orange phenotype observed in *ARF2* VIGS *SlERF.D7-OE#3* line fruits. Furthermore, *SlARF2A/B* VIGS fruits exhibited maximum fruit firmness followed by *SlARF2B* and *SlARF2A*, thereby indicating that a noticeable delay in ripening of VIGS fruits was due to reduced transcript accumulation of *SlARF2* orthologs (Fig. 11). Further, *ARF2A/B* VIGS fruits displayed the lowest ripening index, followed by the *SlARF2B* and *SlARF2A* silenced fruits, respectively (Fig. 11). These results indicate that SlERF.D7 acts upstream of SlARF2 orthologs and performs its ripening-related function by directly regulating them.

**Figure 11.**
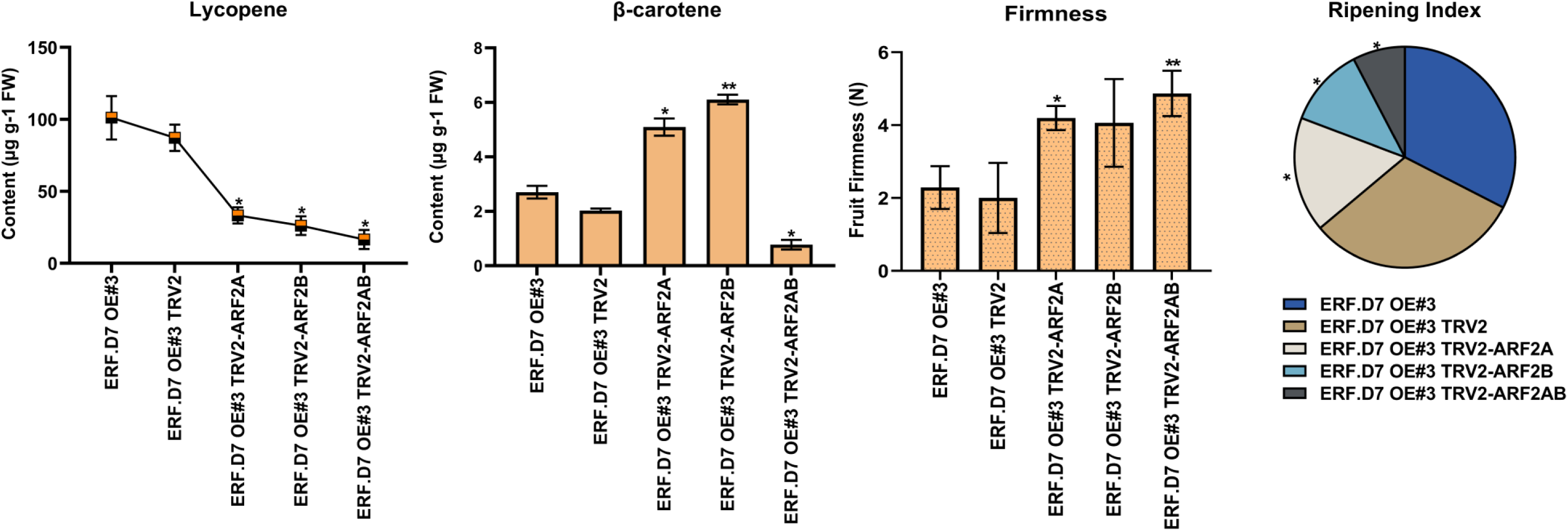
Virus-induced gene silencing of *SlARF2* orthologs in *SlERF.D7* OE transgenic fruits. Assessment of lycopene, β-carotene, fruit firmness levels and ripening index in *SlERF.D7* OE, *SlERF.D7* OE TRV (control), and the *ARF2A*, *ARF2B* and *ARF2AB* VIGS-silenced fruits. Error bars represent ±SD of three biological replicates. Asterisks indicate the statistical significance using Student’s t-test: *, 0.01 < P-value < 0.05; **, 0.001 < P-value < 0.01

## Discussion

### *SlERF.D7*, a ripening-associated gene, encodes a nuclear-localized transcriptional activator

Tomato fruit ripening is a multilayer perception process in which different hormonal inputs operate in parallel or at various levels to define an output transcriptome. For years, the center stage of this ripening process has been occupied by ethylene (Giovannoni, 2004). However, given a large number of ERFs are present in fruit species and the fact that ripening-related roles of many of these are yet to be elucidated, the complete molecular circuitry by which ethylene regulates the ripening-associated genes is not entirely understood (Solano *et al*., 1998; Pirrello *et al*., 2012; Liu *et al*., 2016; Srivastava and Kumar, 2019). For the last two decades, patterns of mRNA accumulation of tomato ERF gene family members during ripening have been under scrutiny. Among the 77 ERFs characterized in tomato, the expression dynamics of several ERFs have demonstrated a classic ripening-associated behavior with an increase in their transcript levels at the onset of ripening, peaking at 5d post breaker, followed by a subsequent decline at late ripening stages (Sharma *et al*., 2010; Liu *et al*., 2016). So far, a limited number of ERFs have been functionally validated for their role in regulating tomato fruit ripening (Liu *et al*., 2016). In this regard, the current study presents several lines of evidence substantiating the physiological significance of SlERF.D7 as an active ripening modulator which corroborates with the well-documented involvement of ERFs in tomato fruit ripening (Srivastava et al., 2019). The presence of ethylene-responsive elements in the promoter region of *SlERF.D7* and its direct regulation by ethylene forms the first line of evidence on this subject. The comprehensive expression profiling of *SlERF.D7* during different stages of fruits in wild-type, RIN silenced genotypes and ripening impaired *rin* and *nor* mutants further establish a strong positive correlation between SlERF.D7 and tomato fruit ripening in multiple genetic backgrounds. A strong transactivation potential in yeast coupled with exclusive nuclear localization, SlERF.D7 protein displays consistency in being designated as a transcriptional factor, as shown previously for the key ripening regulator RIN protein (Ito *et al*., 2008). Although the mRNA abundance of *SlERF.D7* is dramatically repressed in *rin* mutant fruits in parallel with the presence of CArG box in its promoter region, the present research indicates that SlERF.D7 and RIN do not seem to operate in the same regulatory network and its ripening-associate transcription is possibly controlled by some other ripening regulator.

### SlERF.D7 influences all the major facets of fruit ripening

To functionally validate the importance of SlERF.D7 in the ripening process, we generated stable overexpression and RNAi transgenic lines of *SlERF.D7* using a fruit ripening-specific *RIP1* promoter (Agarwal *et al*., 2016). *RIP1* promoter was used in this study to avoid any pleiotropic non-ripening effects in the *SlERF.D7* transgenic lines. The altered phenotypes associated with over and underexpression of *SlERF.D7,* such as ethylene production and fruit pigmentation, signify that this transcription factor broadly impacts the fruit ripening process. As per the fact that a respiratory burst coupled with an increase in ethylene production at the onset of ripening is a hallmark of the climacteric fruits, such as tomato, measurements of ethylene emission in OE and RNAi fruits were in synchronization with the observed delay in ripening of *SlERF.D7* RNAi fruits and an early ripening in the fruits of *SlERF.D7* OE lines (Alexander and Grierson, 2002; Giovannoni, 2004). The paramount role of ACC synthase and ACC oxidase genes in mediating the fruit transition from System 1 (auto-inhibitory) to System 2 (auto-catalytic) ethylene production is well known. Tomato *ACS1* and *ACS6* have shown to mediate System 1 ethylene production in immature fruit, whereas the induction of *ACS2*, *ACS4*, *ACO1,* and *ACO4* genes brings about the characteristic sharp increase in ethylene biosynthesis in System 2 (Nakatsuka *et al*., 1998; Barry *et al*., 2000; Barry and Giovannoni, 2007). The suppression of these genes has been found to inhibit tomato fruit ripening (Hamilton *et al*., 1990; Oeller Paul W. *et al*., 1991; Lincoln *et al*., 1993; Nakatsuka *et al*., 1998; Barry *et al*., 2000). A disturbed expression pattern of *SlACO1*, *SlACS2,* and *SlACS4* genes in *SlERF.D7* transgenic fruits is seemingly responsible for the altered ethylene levels observed in these fruits. Strikingly, exogenous ethylene treatment to compensate for low ethylene levels observed in *SlERF.D7* RNAi fruits failed to restore the normal ripening process. This impairment of *SlERF.D7* RNAi fruits to activate autocatalytic ethylene production could result from the inhibited transcript levels of ethylene receptor genes, *SlETR2*, *SlETR3*, *SlETR4,* and *SlETR5.* A critical evaluation of the significance of the *SlETR3* (*NR*) receptor in regulating ethylene biosynthesis and carotenoid accumulation has already been well-documented (Tieman *et al*., 2000). However, targeting a single ethylene receptor to achieve a delay in fruit ripening has been reported to result in compensatory over-expression dynamics of other ethylene receptor genes (Tieman *et al*., 2000). Thus, the inhibition of four ethylene receptors could account for a considerable loss of ability to perceive and channel the ethylene responses to downstream targets in the RNAi fruits. The inhibited ethylene signalling was validated by the downregulation of EIN2 and EIL3 transcripts, the positive regulators of ethylene responses, in *SlERF.D7* RNAi fruits, which may explain the defective ripening phenotype. Downstream of EILs, the signal branches to ERFs, manipulation of which can result in alterations specific to color, flavor, and texture during ripening (Liu *et al*., 2015, 2016; Xie *et al*., 2016). Notably, a report on the functional significance of SlERF6 in mediating carotenoid accumulation during ripening has already been published (Lee *et al*., 2012). Another APETALA2/ERF gene family member, SlAP2a, has negatively correlated to tomato fruit ripening (Chung *et al*., 2010; Karlova *et al*., 2011). Also, downregulation of another ERF, SlERF.B3, via the dominant repressor technology has been shown to cause a delay in tomato ripening (Liu *et al*., 2014). On analyzing the expression profiles of major ERFs, *SlERF.D7* transgenic fruits displayed dramatic modulations in transcript accumulation of a high number of ERFs consistent with alterations in ethylene production and observed ripening phenotypes. A disparity in mRNA levels of *SlERF.E1* in SlERF.D7#OE and SlERF.D7#RNAi lines is interesting and needs further investigation to validate if it is upregulated due to fluctuations in the transcript abundance of *SlARF2A* and *SlARF2B* (Hao *et al*., 2015).

The enhanced lycopene accumulation during tomato fruit ripening is promoted by an up-regulation of *PSY1* and *PDS* and concomitant repression of lycopene cyclases transcripts. Contrastingly, inhibition in the expression of *PSY1* and *PDS* and the enhanced expression of *β-LYC* and *CYC-β* cyclases promote the conversion of lycopene to β-carotene in tomato fruits (Giuliano *et al*., 1993; Fray and Grierson, 1993; Pecker *et al*., 1996; Ronen *et al*., 2000; Fantini *et al*., 2013; Liu *et al*., 2014). Compared with the wild-type, lycopene to β-carotene ratio in transgenic fruits is significantly modified, conferring a deep red color to *SlERF.D7* OE fruits whereas imparting an orange coloration to *SlERF.D7* RNAi fruits at Br+10 stage. This off-balance is probably a result of modulations in transcript levels of *SlPDS*, *SlPSY1,* and *SlCYCB* genes. The incapacity of *SlERF.D7* RNAi fruits to reach a red color could also be attributed to low levels of ethylene production as the rate-limiting enzyme of carotenoid biosynthesis, *SlPSY1*, is known to be ethylene inducible (Fraser *et al*., 1994). Apart from severe impairment of ethylene biosynthetic genes, the effect of *SlERF.D7* RNAi on the transcript levels of carotenoid pathway genes may also result from the down-regulation of the ripening master regulators, such as *RIN*, *NOR*, and *CNR*. These TFs are known to impact carotenoid accumulation in tomato fruits (Alba *et al*., 2005; Manning *et al*., 2006; Vrebalov *et al*., 2009; Osorio *et al*., 2011). In addition, the enhanced expression of *SlLCYB* and *SlCYCB*, in *SlERF.D7* RNAi fruits perfectly explains the changes in carotenoid content and pigmentation observed in these fruits.

The impact of altered transcript levels of *SlERF.D7* on fruit firmness was evident from the texture analysis of *SlERF.D7* OE and RNAi transgenic lines. Numerous studies have elucidated the importance of early cell wall disintegration as one of the biggest challenges responsible for the post-harvest deterioration of tomato fruit (Meli *et al*., 2010; Yang *et al*., 2017). Although PG2a, PL, and PME enzymes are considered the major contributors to the cell wall degradation process, no significant fluctuations were observed in fruit texture in knockout fruits of *PG2a* (Hall *et al*., 1993; Grierson *et al*., 1993; Yang *et al*., 2017). However, CRISPR-generated mutation in *PL* resulted in firmer fruits with increased shelf life (Li *et al*., 2019). The enhanced softening phenotype of *SlERF.D7* OE fruits is in line with the elevated transcript levels of both *SlPG2a* and *SlPL* at late-breaker stages. It is widely accepted that RIN has hundreds of target genes, including those involved in pathways regulating cell wall loosening, such as *PG2a* (Fujisawa *et al*., 2011, 2013; Zhong *et al*., 2013). We speculate that the overexpression of RIN in *SlERF.D7* OE fruits may also contribute to the early fruit softening phenotype observed in these fruits. Likewise, a significant reduction in mRNA accumulation of *SlPL* and *SlPG2a* combined with the decreased levels of *RIN* in *SlERF.D7* RNAi fruits impeccably synchronizes with fruit firmness studies and the phenotype associated with down-regulation of *SlERF.D7*.

### SlERF.D7 regulates the expression dynamics of major ripening regulators

Tomato genetics and genomics resources are very well developed due to detailed and informative research on pleiotropic mutants such as *rin*, *nor,* and *cnr* (*colorless non-ripening*) (Barry and Giovannoni, 2007; Giovannoni, 2004; Manning et al., 2006; Vrebalov et al., 2002). Inhibition of these genes dramatically and irreversibly impairs the ripening signalling cascade and therefore are termed as master regulators of fruit ripening. Interestingly, *SlERF.D7* RNAi lines fruits also display analogous though milder ripening defects as that of *rin* and *nor* mutant fruits, such as low ethylene emission, disturbed carotenoid accumulation, with altered fruit levels firmness. At the genetic level, ripening-specific silencing of *SlERF.D7* exhibits strong repression of *RIN*, *NOR,* and *CNR* genes at late-breaker stages, plausibly contributing to the incompetency of the RNAi fruits to ripen normally. Apart from RIN, other MADS-domain containing transcription factors such as FRUITFUL1 (FUL1), FRUITFUL2 (FUL2), and AGAMOUS-like1 (TAGL1) play critical roles in fruit ripening regulation by forming multimeric protein complexes with RIN. Suppression of these genes substantially inhibits ripening by blocking ethylene biosynthesis and decreases carotenoid accumulation (Leseberg *et al*., 2008; Vrebalov *et al*., 2009; Itkin *et al*., 2009; Martel *et al*., 2011; Bemer *et al*., 2012). Previous reports on the down-regulation of *TAGL1* have shown to yield a yellow-orange pigmented fruit phenotype accompanied by a drastic decrease in levels of ethylene (Vrebalov *et al*., 2009; Itkin *et al*., 2009). In addition, *FUL1* and *FUL2* are functionally redundant in controlling the expression module of pigment-producing genes during ripening (Bemer *et al*., 2012). Interestingly, the fruit phenotypes observed in *FUL1*, *FUL2,* and *TAGL1* suppression lines are similar to the orange-colored *SlERF.D7* RNAi silenced fruits. Correspondingly, the transcript abundance of these genes in *SlERF.D7* RNAi fruits displays a downward expression profile, keeping in line with the observed phenotype of these fruits. Taken together, *SlERF.D7* emerges as a critical networking component that is transcriptionally wired into a complex regulatory interplay involving several MADS-box proteins essential for ripening to occur.

### SlERF.D7 integrates ethylene and auxin signal transduction pathways via the activation of *SlARF2* orthologs during ripening

Ethylene and auxin signalling components have previously been reported to be arranged and integrated into ways that can modulate the state and output of numerous plant developmental processes. However, precise signal integrators responsible for these interactions remain poorly understood despite a well-defined networking system. There is overwhelming evidence that reinforces ethylene-auxin complex interactions at the molecular level. It has been shown that mutations in auxin signalling display abnormal ethylene responses (Pickett *et al*., 1990*b*). In addition, a mutual regulation, by auxin and ethylene, of genes involved in the biosynthetic machinery of both the hormones has also been reported (Muday *et al*., 2012). In this regard, ERF1 has been designated as a mediator between ethylene and auxin biosynthesis via regulating ASA1 expression in Arabidopsis during root elongation (Mao *et al*., 2016). Likewise, auxin biosynthesis during lateral root formation is directly impacted by AtERF109 (Cai *et al*., 2014). Recently, SlERF.B3 has been reported to directly target SlIAA27 in mediating root length in tomato (Liu *et al*., 2018). Previously, we have also demonstrated the role of SlGH3-2, a ripening-induced IAA-amido synthetase, in regulating fruit ripening aspects by controlling ethylene biosynthesis and auxin homeostasis (SravanKumar et al., 2018). This study’s detailed functional analysis combined with various bioinformatic approaches has rendered SlERF.D7 an active ripening regulator. To add to the complexity, we found that SlERF.D7 functions as an integrator in the interplay between ethylene and auxin through the regulation of ARF2 orthologs, members of the tomato ARF family of transcriptional regulators. The transcriptional regulation of *SlARF2* by both auxin and ethylene has already been reported (Hao *et al*., 2015; Breitel *et al*., 2016*b*). Furthermore, SlERF.D7 positively mediates ethylene and auxin sensitivity, and its silencing results in modifications in ethylene levels, alterations in pigment accumulation, and a decrease in fruit firmness, reminiscent of the phenotypes of the tomato fruits under-expressing *ARF2* orthologs (Hao *et al*., 2015; Breitel *et al*., 2016*b*). Apart from the resemblance in the fruit phenotypes, a slight increase in accumulation of IAA levels in *SlERF.D7* RNAi fruits, in conjunction with significant alterations in the expression of *SlARF2A* and *SlARF2B* in the silenced lines, indicate these genes to be the putative target of SlERF.D7. The presence of ERE elements in the promoters of *SlARF2A* and *SlARF2B* further supported our notion. The heterologous and *in planta* confirmation of SlERF.D7 binding to SlARF2A and SlARF2B promoters in yeast and *N. benthamiana*, respectively, validated this hypothesis. Consistent with the idea of SlARF2 being a direct target of SlERF.D7 in Y1H and electrophoretic mobility shift experiments, subsequent transactivation assays using GUS and GFP reporter systems in *N. benthamiana* evidently demonstrated a direct association between the two classes of transcription factors. Another key piece of evidence further substantiating the activation of *ARF2* promoters by SlERF.D7 is obtained from the VIGS assay of single and chimeric knock-down of *ARF2A/B* in *SlERF.D7* OE lines. The ripening phenotypes exhibited by *SlARF2A*, *SlARF2B* and *SlARF2A/B* VIGS fruits are in agreement with the similarity of phenotypes observed in *SlERF.D7* RNAi and *SlARF2* silenced fruits (Hao *et al*., 2015; Breitel *et al*., 2016*b*). Altogether, the data reveals that SlERF.D7 acts upstream of SlARF2 transcription factors and impacts fruit ripening via their transcriptional activation.

In summary, our results underpin a prototype in which the ripening-induced ethylene response factor, SlERF.D7, emerges as a critical positive regulator of fruit ripening. SlERF.D7 amalgamates auxin and ethylene signalling pathways via regulating the transcript accumulation of SlARF2 orthologs (directly) and other key ripening regulators (unknown mechanism) in tomato (Fig 12). Silencing of *SlERF.D7* culminates in down-regulation of *SlARF2A* and *SlARF2B* genes, negatively impacting tomato fruit ripening. The SlERF.D7-ARF2A/B association seems critical for fine-tuning ethylene and auxins aspects, such as pigment accumulation and shelf-life during fruit ripening. Nonetheless, further research is warranted to delineate the molecular mechanisms underlying SlERF.D7 controlled ripening regulation and validate if SlERF.D7 directly activates the transcription of ripening master regulators such as RIN, NOR, and CNR by binding to their promoters. Besides identifying the transcription factor responsible for its activation during ripening, SlERF.D7 physical binding to ERE elements of the promoters of ARF2 orthologs and other transcriptionally affected genes in this study also remains to be ascertained.

**Figure 12.**
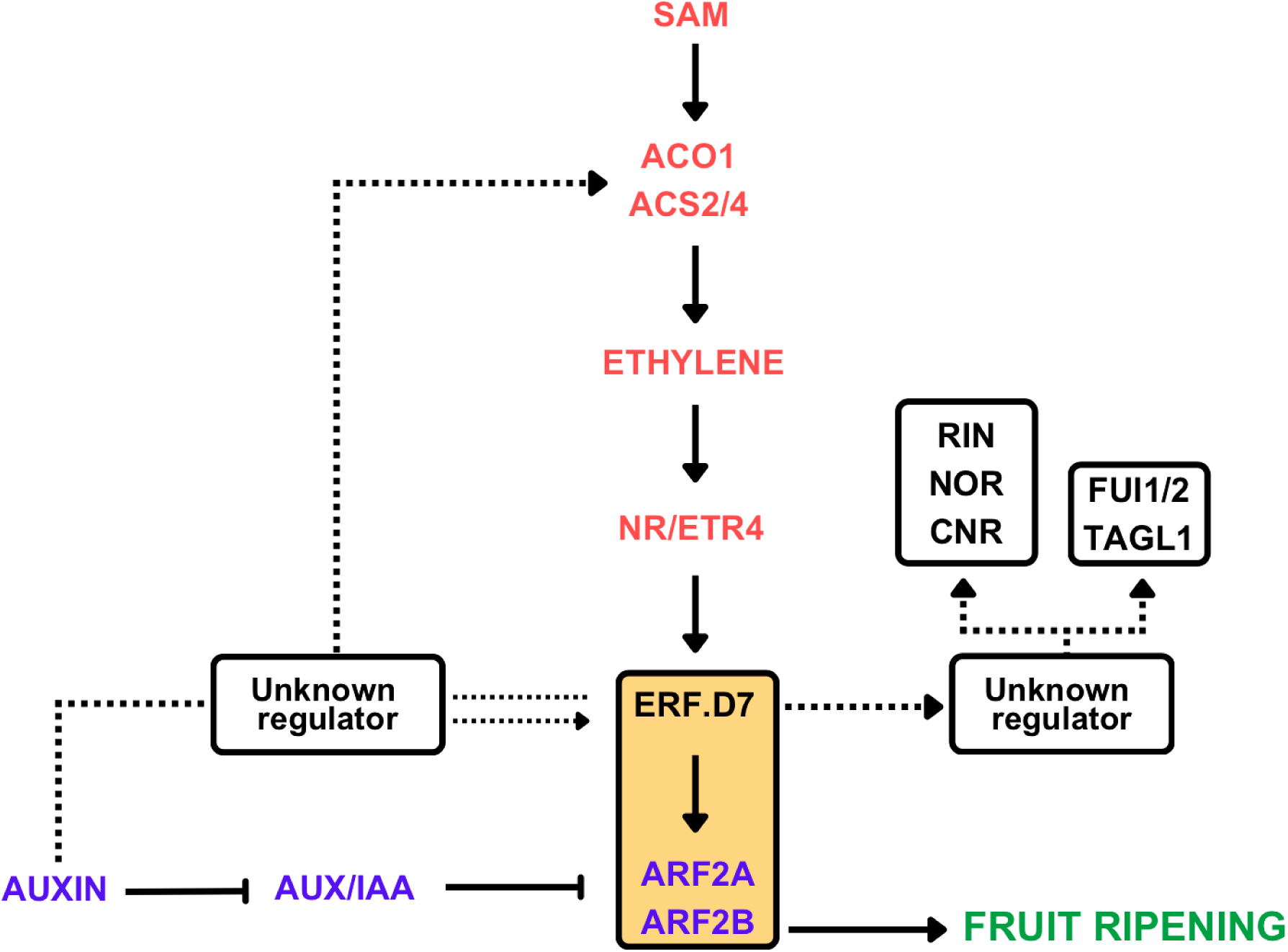
A general regulatory model depicting the role of SlERF.D7 in controlling fruit ripening. *SlERF.D7* positively regulates tomato fruit ripening via modulating the transcript profiles of key ripening regulators such as RIN, NOR, and CNR associated with the expression of important ethylene (*SlACO1*, *SlACS2/4*) and carotenoid (*SlPSY*, *SlPDS*) biosynthesis genes as well as other ripening metabolic pathways. To add to the complexity, *SlERFD7* also targets the transcript levels of TAGL and FUL1/FUL2 via an unknown regulator. Furthermore, SlERF.D7 works in ethylene and auxin-dependent manner and activates the expression of *SlARF2A* and *SlARF2B*; thus, down-regulation of SlERF.D7 displays a severe impairment of ARF2 orthologs. Functional and physiological dissection of the mode of action of ARF2 states that it positively correlates with the fruit ripening process. The dotted lines represent the putative regulatory circuits indicating the probability of an unknown regulator controlling the part of the circuit. Taken together, SlERF.D7 emerges as an integrator of ethylene and auxin signalling pathways working synergistically to achieve competency to ripen.

## Supplementary Data

**Table S1** List of primers used in this study.

**Table S2** Segregation analysis of kanamycin resistance in T_2_ progeny of *SlERF.D7 OE* and *SlERF.D7* RNAi transgenic tomato plants.

**Figure S1** DAPI staining of SlERF.D7 in *Nicotiana benthamiana*.

**Figure S2** The expression of *ERF* genes in wild-type and *SlERF.D7 OE* and *SlERF.D7 RNAi* plants.

**Figure S3** The expression of ERF.D clade genes in wild-type and *SlERF.D7 OE* and *SlERF.D7 RNAi* plants.

**Figure S4** The expression of *SlARF* in wild-type and *SlERF.D7 OE* and *SlERF.D7 RNAi* plants.

## Acknowledgements

This work is financially supported by grants received from the Department of Biotechnology (DBT), Government of India. RK Acknowledges the Department of Science and Technology, India, for the INSPIRE-Faculty (IF-LSPA-15) grant. AKS acknowledges DBT grant BT/PR6983/PBD/16/1007/2012. The authors acknowledge the Department of Science and Technology, India, for the Purse Grant. The SAP Grant of the University Grants Commission and FIST grant of DST, India, to the Department of Plant and Molecular Biology for infrastructure support are also acknowledged. There is no conflicts of interest to report.

## Author contributions

Conceptualisation- A.K.S., R.K.; Investigation, validation, methodology and formal analysis- P.G., A.P., U.R., V.S.; Writing original draft-P.G.; Reviewing and editing- A.K.S., R.K.; Supervision- A.K.S., R.K..

## Abbreviations

ERF: Ethylene response factor
ARF: Auxin response factor
1-MCP: 1 Methylcyclopropane
PCIB: p-Chlorophenoxyisobutyric acid
IAA: Indole-3-acetic acid
VIGS: Virus induced gene silencing

## Accession Numbers

The sequences of genes used for the qPCR can be found at the website (http://solgenomics.net/) under the following solyc numbers: *Sl-ERF*.*A3* (Solyc06g063070), *SlERF*.*B1* (Solyc05g052040), *SlERF*.*B2* (Solyc02g 077360), *SlERF*.*B3* (Solyc05g052030), *SlERF*.*C1* (Solyc05g051200), *SlERF*.*D1* (Solyc04g051360), *SlERF*.*D2* (Solyc12g056590), *SlERF*.*D3* (Solyc01g108240), *SlERF*.*D4* (Solyc10g050970), *SlERF.D5* (Solyc04g012050), *SlERF.D6* (Solyc04g071770), *SlERF.D7* (Solyc03g118190), *SlERF.D8* (Solyc12g042210), *SlERF.D9* (Solyc06g068830), *Sl-ERF*.*E1* (Solyc09g075420), *Sl-ERF*.*E2* (Solyc09g089930), *SlERF*.*E2* (Solyc06g063070), *SlERF*.*E4* (Solyc01g 065980), *PSY1* (Solyc03g031860), *PDS* (Solyc03g123760), *ZDS* (Solyc01g097810), *βLCY1* (Solyc04g040190), *βLCY2* (Solyc10g079480), *CYCβ* (Solyc06g074240), *ACS2* (Solyc01g095080), *ACS4* (Solyc05g050010), *ACO1* (Solyc07g049530), *E4* (Solyc03g111720), *E8* (Solyc09g089580), *PG2a* (Solyc10g080210), *R IN* (Solyc05g012020), *CNR* (Solyc02g077920), *NOR* (Solyc10g006880), *TAGL 1* (Solyc07g055920), *AP2a* (Solyc03g044300), *EIN2* (Solyc09g007870), *EIL2* (Solyc01g009170), *EIL3* (Solyc01g096810), *ETR2* (Solyc07g056580), *ETR3* (So lyc09g075440), *ETR4* (Solyc06g053710), *ETR5* (Solyc11g006180),*FUL1* (Soly c06g069430), *FUL2* (Solyc03g114830), *ACO2* (Solyc12g005940), *ACO3* (Soly c07g049550), *ACO4* (Solyc02g081190). SAUR (Solyc09g007970.1.1), *ARF2A* (Solyc03g118290), *ARF2B* (Solyc12g042070).

